# Calcium and actomyosin pulses drive tricellular zipping to remodel epithelial cell-cell junctions and facilitate cell packing

**DOI:** 10.64898/2026.10.30.763104

**Authors:** Babli Adhikary, Xinran O. Zhao, Ashley Shahi, Andrea G. Laboy-Figueroa, Michaela P. Purvis, Katherine K. Tjoelker, Tyler R. Huycke, Shiladitya Banerjee, Adam C. Martin, Ann L. Miller

**Affiliations:** Department of Molecular, Cellular, and Developmental Biology, University of Michigan, Ann Arbor, MI, USA; Department of Biology, Massachusetts Institute of Technology, Cambridge, MA, USA; Cellular and Molecular Biology Graduate Program, University of Michigan, Ann Arbor, MI, USA; School of Physics, Georgia Institute of Technology, Atlanta, GA, USA

**Keywords:** Tricellular zipping, cell packing, calcium flashes, actomyosin, pulsatile contraction, cell-cell junctions, junction remodeling, vertex remodeling, tricellular junction, live microscopy, computational modeling, *Xenopus laevis*, *Drosophila*

## Abstract

Epithelial morphogenesis requires coordinated changes in cell geometry, neighbor cell arrangement, and cell packing to generate functional tissue architectures. However, the spatiotemporal regulation of upstream signals that drive local junctional remodeling in highly proliferative developing epithelial tissues remains unclear. Here, we characterize a new junction remodeling process, which we call ‘tricellular zipping’. It resolves long, thin cell extensions generated by cell-division forces that terminate at transient higher-order vertices. We show that tricellular zipping facilitates epithelial cell packing in blastula-stage *Xenopus laevis* embryos through asymmetric displacement of a tricellular vertex and formation of a *de novo* bicellular interface. Similarly, we observe tricellular zipping in developing mouse embryos. Combining modeling and live imaging approaches, we find that displacement of the remodeling vertex depends on actomyosin-dependent asymmetric force generation and on enrichment of adaptor proteins coupling actomyosin to junctions, which enables effective force transmission. In contrast, the stationary vertex remains close to mechanical equilibrium and therefore undergoes no net displacement. Furthermore, we find that calcium flashes drive the localized, pulsatile actomyosin contractility required for efficient tricellular zipping. Extending our observations, we show that disrupting transient calcium signaling also affects cell packing in *Drosophila* embryos, thereby impairing mesoderm internalization. Overall, by combining multidisciplinary and multisystem approaches, we uncover a new mode of junctional remodeling and highlight the importance of spatiotemporally regulated calcium signaling in driving local junctional rearrangements to facilitate morphogenesis.

## Introduction

Morphogenesis and organogenesis in developing organisms rely on both the structural order and active mechanical machinery provided by polarized epithelial tissues. The spatial organization and geometry of cells within these tissues is commonly referred to as epithelial cell packing. While textbook depictions often present epithelia as highly ordered hexagonal arrays of cells, *in vivo* these tissues exhibit dynamic, context-dependent, tightly regulated rearrangements in cell packing^1, 2^. Physical and computational models suggest that epithelia can transition between a solid-like ‘jammed’ state, in which cells rarely exchange neighbors, and a fluid-like ‘unjammed’ state characterized by frequent cell rearrangements and changes in cell geometry^1, 3–5^. Many morphogenetic and homeostatic processes involve phase transitions in packing dynamics, with increased cell rearrangements driving cell packing transitions that ultimately give rise to functional tissue architectures^6^. For example, changes in epithelial packing are critical for developmental processes including body axis elongation^7–9^, gastrulation^10, 11^, left-right axis establishment^12^, gland and duct formation^1, 13^, and epithelial to mesenchymal transition^14^. Furthermore, cell division^15–21^, cell extrusion^22–24^, and apical constriction^25–27^ result in strain of the surrounding tissues that necessitate local rearrangements in order to relax.

Epithelial packing is facilitated by remodeling of specialized cell-cell junctions that mediate adhesion and barrier function along bicellular interfaces and at tricellular and multicellular vertices. These junctions include tight junctions (TJs) (or septate junctions (SJs) in invertebrates), which seal the paracellular space and form a selective barrier^28–33^ as well as adherens junctions (AJs)^34–36^ and desmosomes^37–39^, which provide adhesion and mechanical coupling between neighboring cells. Additionally, specialized proteins localize at tricellular junctions (TCJs) to regulate barrier function and adhesion at these cellular vertices^40–42^. TJs and AJs are associated with the actomyosin cytoskeleton to maintain tissue architecture and transmit forces across the tissue^43, 44^. Epithelial tissues must balance maintenance of adhesion and barrier function with the need to dynamically remodel their junctional complexes to enable changes in cell shape and tissue packing^45, 46^.

Several cell-cell junction rearrangement processes have been characterized including planar cell polarized junctional contraction^47–51^, asymmetric vertex movement^50, 52, 53^, post-cytokinesis remodeling^15, 16, 19–21^, and multicellular zippering^54, 55^. Cell intercalation, observed in diverse biological contexts such as *Drosophila* germband extension^56^ and *Xenopus* neurulation^57^, involves T1 transitions: (1) shortening of an existing junction, (2) formation of a transient multicellular vertex involving four (T1 configuration) or more (rosette configuration) cells^58^, and (3) elongation of a new cell-cell junction along the orthogonal axis^51, 56^.

Junction shortening can also occur via asymmetric vertex sliding, in which local differences in actomyosin contractility, cadherin-mediated adhesion, or junctional stiffness bias the movement of one vertex, although the underlying mechanisms vary by context. For example, during *Drosophila* germband extension, oscillatory Myosin II and E-cadherin dynamics at the motile vertex correlate with cell area changes and promote sliding^50^. Local heterogeneity along the junction also biases vertex motion in Caco-2 cells, where RhoA-dependent E-cadherin accumulation stabilizes the less motile vertex^53^, and in *Xenopus* embryos, where Cadherin-3 clustering promotes local stiffening near the active (motile) vertex^52^.

The formation and elongation of new cell-cell junctions is also regulated by coordinated cytoskeletal and adhesive dynamics. During *Drosophila* intercalation, polarity proteins and TCJ proteins regulate E-cadherin and Myosin II dynamics at elongating cell contacts^59–62^. Following cytokinesis in epithelia, new bicellular junctions (BCJs) form and drive local changes in packing order. In the *Drosophila* embryo, formation of a daughter-daughter interface following cytokinesis is driven by Myosin II activity in neighboring cells and Arp2/3-mediated actin assembly in the dividing cell, strengthening adhesion at the newly forming BCJ^16, 63^. Whether similar mechanisms operate during cell intercalation or post-cytokinesis remodeling in vertebrate epithelia remains unclear.

A central theme of epithelial junction remodeling is the interplay between actomyosin dynamics and junctional complexes, but the upstream signaling mechanisms that drive junction remodeling in different developmental contexts remains unclear. In particular, little is known about how junction remodeling coordinates with rapid cell divisions to establish a well packed epithelium in developing vertebrate embryos. Here, we identify a calcium-dependent junction remodeling process that responds to morphogenetic events that challenge epithelial packing. In this process, which we term ‘tricellular zipping’, a long, thin cell extension that forms following cytokinesis is resolved as two neighboring cells zip together through elongation of a new BCJ, thereby repositioning the tricellular vertex and facilitating cell packing during the *Xenopus* blastula-to-gastrula transition. We observe similar tricellular zipping in developing mouse embryos. Live microscopy in *Xenopus* embryos combined with computational modeling show that unidirectional movement of the remodeling vertex requires asymmetric force generation and transmission, whereas the stationary vertex remains close to mechanical equilibrium. Dominant negative and pharmacological perturbations, live imaging with a genetically-encoded calcium indicator, calcium/F-actin cross-correlation analysis, local calcium uncaging, and simulations reveal that tricellular zipping is dependent on localized calcium flashes that drive pulsatile actomyosin contractility and accelerate vertex movement. Furthermore, we show that calcium signaling also regulates epithelial cell packing during presumptive mesoderm invagination in *Drosophila* embryos. Together, these findings establish spatiotemporally regulated calcium signaling as a conserved mechanism linking junctional remodeling to epithelial cell packing during morphogenesis.

## Results

### Tricellular zipping facilitates hexagonal epithelial cell packing during the blastula-to-gastrula transition in *Xenopus* embryos

During gastrulation, the outer surface of *Xenopus laevis* embryos is covered with a polarized epithelium with apical cell-cell junctions facing the external environment, making them an ideal system to investigate changes in epithelial packing and junctional reorganization. To determine whether packing order changes as the embryo enters gastrulation, we imaged *Xenopus* embryos beginning at approximately 6 hours post-fertilization, corresponding to Nieuwkoop and Faber late stage 8 (blastula) through stage 10 (gastrula)^64^ **(Fig. 1A and Video S1)**. As embryos underwent the blastula-to-gastrula transition, cells changed from an irregular morphology with long, thin extensions to a more regular, hexagonally packed organization **(Fig. 1A)**. Consistent with this transition, mean cell circularity increased, and the cell shape index decreased over time **(Fig. 1B and S1A, B)**, highlighting the emergence of a more regularly packed epithelium.

**Figure 1.**
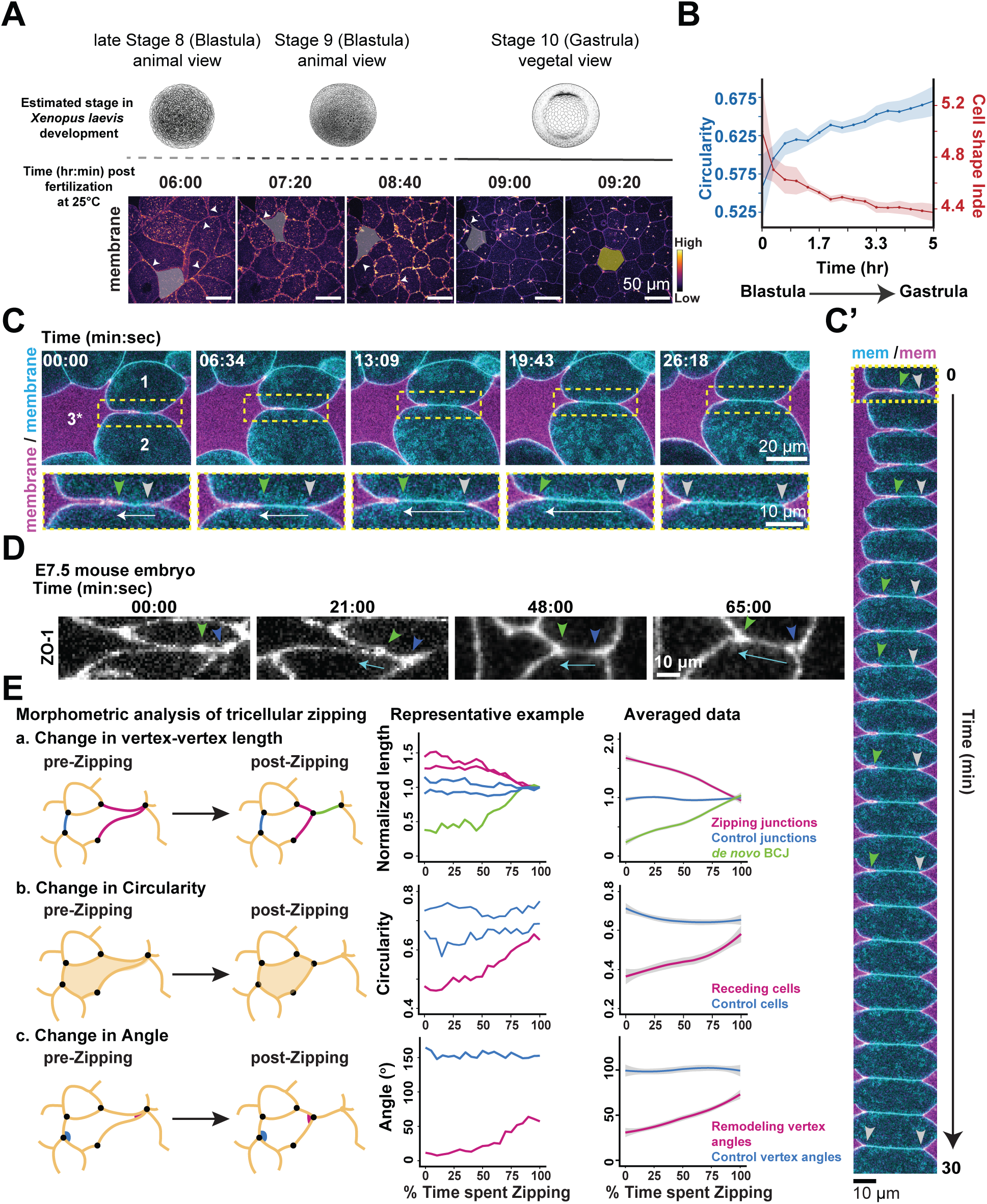
Tricellular zipping facilitates epithelial cell packing during the blastula-to-gastrula transition in *Xenopus* embryos. **(A)** Time-lapse confocal images of a *Xenopus laevis* embryo expressing a membrane probe (mem-BFP). Gray overlays highlight irregularly shaped cells with long cellular extensions (white arrowheads), and yellow overlay highlights a cell with more regular, hexagonal packing. Embryonic stage illustrations are from Zahn et al., 2022^86^. Time shown as hr:min post-fertilization. Scale bars, 50 µm. **(B)** Quantification of mean cell circularity (blue) and cell shape index (red) for cells during the blastula-to-gastrula transition. Mean (solid line) ± SEM (shading). *n* = 6 embryos, 3 experiments. **(C)** Time-lapse confocal images of a *Xenopus laevis* embryo mosaically expressing two membrane probes (mem-BFP and mCherry-farnesyl). The long cell extension (yellow box, enlarged below) is resolved via lengthening of the *de novo* bicellular interface (between cells labeled 1 and 2, zipping cells). Remodeling vertex (green arrowheads), stationary vertex (gray arrowheads), direction of zipping (white arrows), receding cell (3*). Time shown as min:sec from the onset of zipping. Scale bars, 20 µm (upper), 10 µm (lower). **(C′)** Montage generated from the mosaic membrane signal shown in (C). Remodeling vertex (green arrowheads), stationary vertex (gray arrowheads). Scale bar, 10 µm. **(D)** Time-lapse confocal images of tricellular zipping in a ZO-1-GFP E7.5 mouse embryo. Remodeling vertex (green arrowheads), stationary vertex (blue arrowheads), direction of zipping (cyan arrows). Time shown as min:sec from the onset of zipping. Scale bar, 10 µm. **(E)** Quantification of morphometric changes during tricellular zipping. Schematics illustrate the zipping process and the measured parameters. (a) Vertex-to-vertex lengths of zipping junctions (magenta), *de novo* BCJs (green), and control junctions (blue) were measured over time and normalized to their respective initial lengths at the onset of zipping. (b) Circularity was measured for receding cells involved in zipping (magenta) and control cells within the same field of view that did not participate in zipping (blue). (c) The angle of the remodeling vertex (magenta) and control vertices (blue) were measured during zipping. Representative example plots show a single zipping event. Data trend plots show nonlinear regression across multiple zipping events. The normalized time axis represents the percentage of the total time spent zipping. Fitted mean (solid line) ± 95% confidence interval (shading). For (a), *n* = 10 *de novo* BCJs, 17 zipping junctions, and 23 control junctions; 9 embryos; 3 experiments. For (b), *n* = 10 zipping cells and 17 control cells; 7 embryos; 3 experiments. For (c), *n* = 10 remodeling vertices and 14 control vertices; 7 embryos; 3 experiments.

To investigate how these cell extensions are resolved, we generated late blastula-stage *Xenopus laevis* embryos in which the epithelium contained mixed cell populations expressing two differently colored fluorescently tagged membrane probes. We imaged mosaic regions containing cells with long extensions **(Fig. 1C and Video S2)**. We observed that cell extensions terminating at higher-order junctions (where 4 or more cells come together) were resolved as two cells zipped together by lengthening a new bicellular interface, while the cell with extension receded. During this process, the two vertices exhibited asymmetric behavior: the tricellular vertex associated with the receding cell moved unidirectionally away from the other vertex and will therefore be referred to as the ‘remodeling vertex’, while the second vertex did not move significantly and will be referred to as the ‘stationary vertex’ **(Fig. 1C’)**. We visualized the same behavior in embryos labeled with a cell-permeable membrane dye, confirming that it is not an artifact of membrane probe expression **(Fig. S1D)**. Because this vertex remodeling process has not, to our knowledge, been previously documented, we refer to it as ‘tricellular zipping’. Interestingly, we also observed tricellular zipping in the surface endoderm of embryonic day (E)7.5 mouse embryos **(Fig. 1D)**, suggesting that similar junction remodeling also happens in other developing vertebrate embryos. To leverage the ease of experimental accessibility and live imaging in developing *Xenopus* embryos, we further characterized tricellular zipping in *Xenopus*.

We next performed morphometric analysis to better understand the shape and geometry of cells and junctions involved in tricellular zipping. Zipping junctions decreased in length as the remodeling vertex moved, while the *de novo* BCJ steadily increased in length **(Fig. 1Ea)**. The receding cell became more circular as zipping progressed **(Fig. 1Eb)**. In a regularly packed epithelium, the interior vertex angles are typically obtuse (>90°, with a perfect hexagon having interior vertex angles of 120°). Indeed, the control vertex angles are obtuse **(Fig. 1Ec)**. In contrast, the long cell extension creates an extremely acute angle at the higher-order vertex, and as the remodeling vertex moves, this angle gradually becomes more obtuse, indicating a change in the packing geometry of the receding and zipping cells **(Fig. 1Ec)**. The time required to complete tricellular zipping was variable, typically ranging from 30 to 60 min. Of note, zipping duration correlated with the change in cell circularity **(Fig. S1C**), indicating that cells that experience a larger increase in circularity take a longer time to complete zipping.

We also observed a previously-characterized form of asymmetric vertex movement – vertex sliding^50^ – in the *Xenopus* epithelium during the blastula-to-gastrula transition **(Fig. S1E, E’)**. Although both tricellular zipping and vertex sliding involve asymmetric vertex motion and lead to junction elongation, we identified several key differences. First, vertex sliding is defined by the shifting of a vertex along an existing cell-cell contact, whereas in tricellular zipping, vertex movement occurs in concert with membrane convergence to create a *de novo* bicellular interface. Thus, the change in vertex-to-vertex length is smaller for vertex sliding compared with tricellular zipping **(Fig. 1E and S1F)**. Second, the initial cell geometry differs between the two processes: vertex sliding is observed in cells starting with a higher circularity index (>0.6), whereas tricellular zipping occurs in cells starting with a much lower circularity index (0.4) **(Fig. 1E and S1F)**. In addition, the initial remodeling vertex angle is much more acute in tricellular zipping (∼>25°) than in vertex sliding (∼50°) **(Fig. 1E and S1F)**.

Together, these data indicate that tricellular zipping is a vertex remodeling process where two neighboring cells zip together by elongating a new bicellular interface, thereby relocating the tricellular vertex. This process contributes to the transition from irregular cell shapes at blastula stage to a more regularly packed epithelium in gastrula-stage *Xenopus* embryos.

### Tricellular zipping resolves cellular extensions that arise during cell division

We next wanted to determine: what causes the long cellular extensions that are resolved by tricellular zipping to form during early *Xenopus* embryogenesis? During the blastula-to-gastrula transition, embryos undergo frequent cell divisions. We found that during blastula stage, dividing cells pulled neighboring cells in with the cleavage furrow, creating the long, thin extensions **(Fig. 2A, B and Video S3)**. We frequently observed that these cell extensions formed between two daughter cells as the cytokinetic furrow ingressed asymmetrically **(Fig. 2A)** or were pulled from neighboring cells during division **(Fig. 2B)**. To determine the position of the remodeling vertex and stationary vertex with respect to the midbody, we expressed a member of the Centralspindlin complex, MgcRacGAP, as a midbody marker^19^. The midbody was located at the tip of the cell extension, adjacent to the transient, higher-order vertex formed at the end of cytokinesis **(Fig. 2C)**. Tricellular zipping began on average 25.46 ± 3.4 min after cytokinesis completion **(Fig. 2D)**. The remodeling vertex consistently moved away from the midbody with a significantly higher velocity than the stationary vertex, which did not move appreciably relative to the midbody **(Fig. 2E, F)**. A previous study showed that the directionality of furrow ingression changes from cutting apical-to-basal in blastula-stage embryos to progressing basal-to-apical in gastrula-stage embryos^65^. We investigated whether the directionality of tricellular zipping showed apicobasal polarity and found that tricellular zipping progressed from the basolateral to the apical surface **(Fig. 2G)**. Together, these data suggest that tricellular zipping resolves cell extensions formed after cytokinesis and contributes to epithelial packing during early *Xenopus* embryogenesis when frequent cell divisions may challenge epithelial packing order.

**Figure 2.**
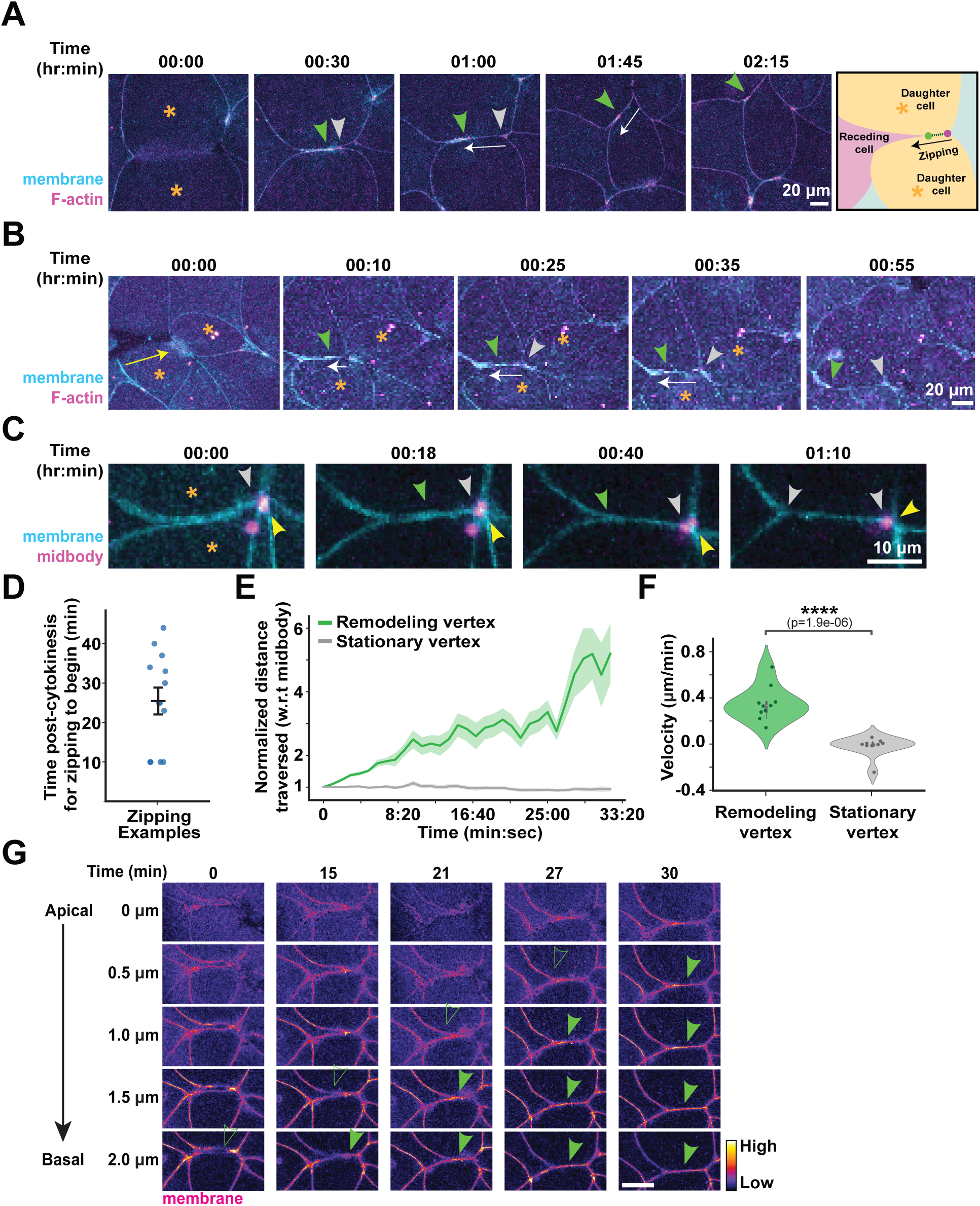
Tricellular zipping resolves cellular extensions that arise due to cell division. (**A-B**) Time-lapse confocal images showing tricellular zipping associated with resolution of cellular extensions generated by cell division (A) and with resolution of cellular extensions generated by pulling forces during cell division (B); yellow arrow shows the directionality of pull by dividing cell). *Xenopus* embryos are expressing probes for membrane (mem-BFP) and F-actin (LifeAct-miRFP703). Daughter cells (orange asterisks), remodeling vertex (green arrowheads), stationary vertex (gray arrowheads), direction of zipping (white arrows). Time shown as hr:min from the onset of zipping. Scale bars, 20 µm. **(C)** Time-lapse confocal images of tricellular zipping in early *Xenopus* embryos expressing a membrane probe (mem-BFP) and a midbody marker (MgcRacGAP-3xGFP). The remodeling vertex, (green arrowheads) moves away from the midbody (yellow arrowheads), whereas the stationary vertex (gray arrowheads) shows minimal displacement relative to the midbody. Time shown as hr:min from the onset of zipping. Scale bar, 10 µm. **(D)** Quantification of the time post-cytokinesis when zipping begins. Mean ± SEM. *n* = 11 zipping events, 7 embryos, 3 experiments. **(E)** Quantification of the normalized distance traveled by the remodeling vertex (green) and stationary vertex (gray) relative to the midbody during tricellular zipping. Solid lines indicate the Mean (solid line) ± SEM (shading). **(F)** Violin plot showing the velocities of remodeling and stationary vertices calculated from (E). For (E) and (F); *n* = 10 remodeling vertices and 10 stationary vertices, 5 embryos, 3 experiments. Statistical significance in (F) was determined using a student *t*-test; *\*\*\*\*p ≤* 0.0001. **(G)** Time-lapse single z-plane confocal images of tricellular zipping in *Xenopus* embryos expressing a membrane probe (mem-BFP; FIRE LUT). Tricellular zipping proceeds in a basal-to-apical direction (closed green arrowheads: zipped to open green arrowheads: zipping in progress). Time shown in min. Scale bar, 10 µm.

### Tension asymmetry and differential force transmission drives displacement of the remodeling vertex

A key feature of tricellular zipping is the asymmetric motion of the two vertices: the remodeling vertex moves unidirectionally to form the *de novo* BCJ, whereas the stationary vertex exhibits negligible movement. To understand what drives this asymmetric vertex motion, we analyzed vertex displacement within a mechanical force-balance framework **(Fig. 3A and Supplementary Information).**

**Figure 3.**
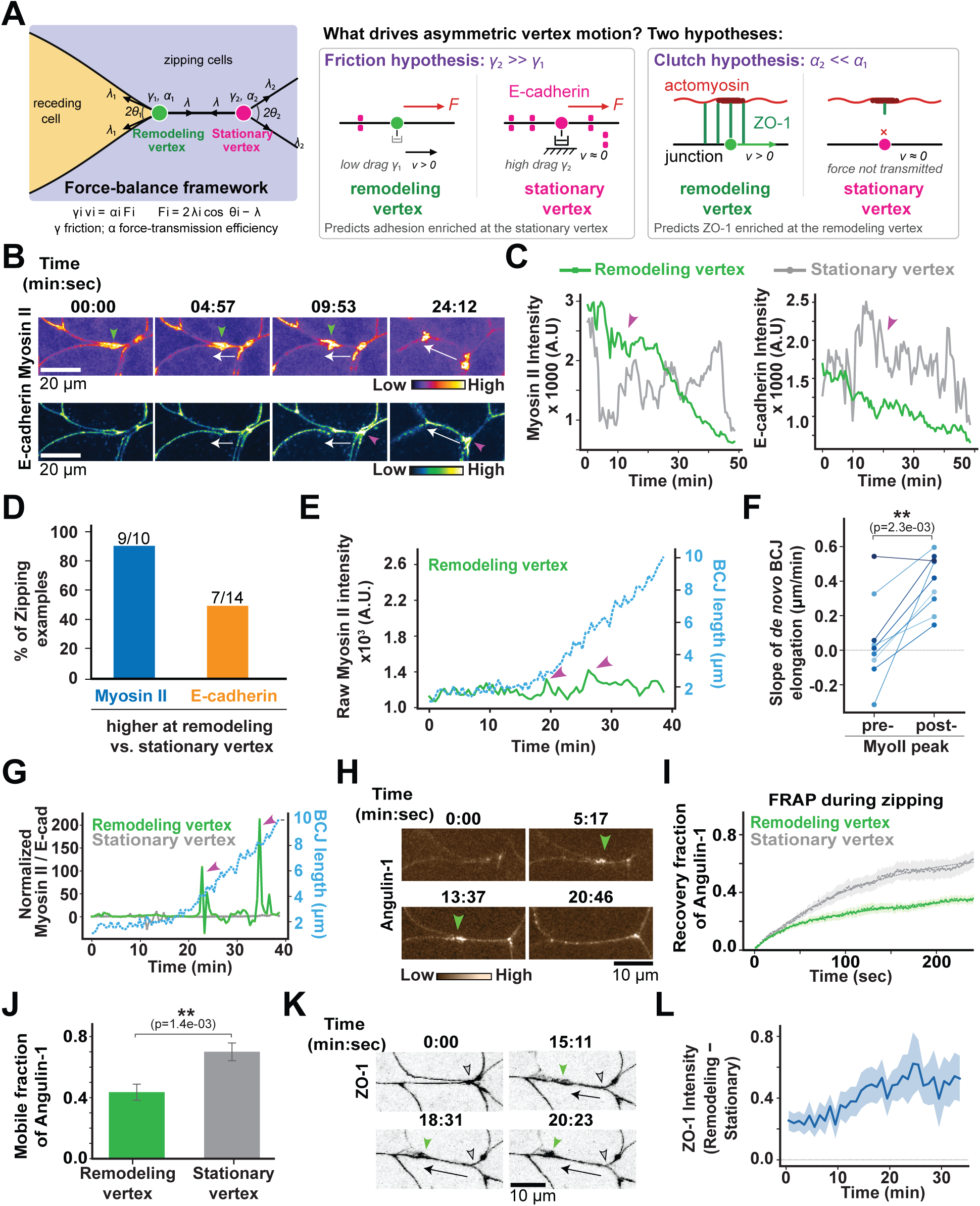
Mechanical forces regulating asymmetric vertex motion. **(A)** Mechanical framework for asymmetric vertex motion. Left: geometry of a zipping event. The receding cell (yellow) forms a thin extension that terminates at the remodeling vertex (green). The BCJ connects it to the stationary vertex (magenta), carrying a tension λ. The shoulder junctions of vertex i carry tension λ_i_ and make half-angle θ_i_ with the zipping axis. Vertex motion is overdamped, so the velocity of vertex i is set by the transmitted net force, γ_i_ v_i_ = α_i_ F_i_, with F_i_ = 2λ_i_ cos θ_i_ − λ, where γ is a friction coefficient and α the efficiency of force transmission. Right: two hypotheses for what drives asymmetric vertex motion. The friction hypothesis proposes that the stationary vertex is anchored by high adhesion-based drag (γ₂ >> γ₁), predicting enrichment of adhesion machinery at the stationary vertex. The clutch hypothesis proposes that the stationary vertex is poorly coupled to actomyosin (α₂ << α₁), so that force is not transmitted to it, predicting enrichment of ZO-1 at the remodeling vertex. **(B)** Time-lapse confocal images of tricellular zipping in *Xenopus* embryos expressing probes for Myosin II (SF9-mCherry; FIRE LUT) and E-cadherin (E-cadherin-3xGFP; Green Fire Blue LUT). Myosin II is enriched at the remodeling vertex (green arrowheads), whereas E-cadherin accumulates at the stationary vertex (magenta arrowheads). The white arrow indicates the direction of zipping. Time shown in min:sec. Scale bars, 20 µm. **(C)** Representative plot showing raw Myosin II and E-cadherin intensity at the remodeling (green) and stationary (gray) vertices during zipping. Myosin II intensity is higher at the remodeling vertex (magenta arrowhead, left) during the early stages of zipping, whereas E-cadherin intensity is higher at the stationary vertex (magenta arrowhead, right). **(D)** Bar graph showing the percentage of zipping events with higher Myosin II or E-cadherin intensity at the remodeling vertex. **(E)** Representative plot showing raw Myosin II intensity at the remodeling vertex, together with the length of the *de novo* BCJ during a zipping event. Magenta arrowheads indicate peaks in the Myosin II intensity. **(F)** Quantification of paired slopes of *de novo* BCJ elongation, before and after a peak in Myosin II intensity (like those shown in E). *n* = 9 zipping events, 7 embryos, 4 experiments. Statistical significance was determined using a paired *t*-test; *\*\*p ≤ 0.01*. **(G)** Representative plot showing the ratio of normalized Myosin II and E-cadherin intensities (Myosin II/E-cadherin) at remodeling (green) and stationary (gray) vertices, together with length of the *de novo* BCJ during a zipping event. Magenta arrowheads indicate peaks in the Myosin II/E-cadherin intensity at the remodeling vertex. **(H)** Time-lapse confocal images of tricellular zipping in *Xenopus* embryos expressing Angulin-1-3xGFP (sepia LUT). Angulin-1 accumulates at the remodeling vertex (green arrowheads) and moves in the direction of zipping. Time shown in min:sec. Scale bar, 10 µm. **(I)** Recovery curves for Angulin-1-3xGFP FRAP at remodeling (green) and stationary (gray) vertices, fitted with a single-exponential nonlinear regression (dashed lines). Shaded regions indicate SEM. *n* = 5 experiments; 21 remodeling vertices from 10 embryos; 20 stationary vertices from 9 embryos. **(J)** Mobile fractions calculated from (I). Error bars indicate SEM. Statistical significance was determined using Mann-Whitney U test; *\*\*p ≤ 0.01*. **(K)** Time-lapse confocal images of tricellular zipping in *Xenopus* embryos expressing TagBFP2-ZO-1. ZO-1 accumulates at the remodeling vertex (green arrowheads) and moves in the direction of zipping (black arrow). Stationary vertex (outlined gray arrowheads). Time shown in min:sec. Scale bar, 10 µm. **(L)** Quantification of the difference in ZO-1 intensity between the remodeling and stationary vertex. Mean (solid line) ± SEM (shading). *n* = 13 remodeling vertices and 13 stationary vertices, 8 embryos, 7 experiments.

Because vertex motion in the epithelium is overdamped, the velocity of a vertex is proportional to the net force acting on it. A vertex moves when the forces exerted by the junctions meeting there do not cancel, while it is stationary when they balance. This framework allows us to ask two distinct questions: What generates the net force that displaces the remodeling vertex? Why does the stationary vertex experience no net force, despite being embedded in the same tension network?

To answer these questions, we develop a mechanical model of tricellular zipping and test two hypotheses. First, we test the *friction hypothesis*, where the stationary vertex is subject to active force but is held in place by a high local resistance to motion, such as adhesion-based friction. As a result, the friction coefficient γ is expected to be much larger at the stationary vertex than at the remodeling vertex. The second, *clutch hypothesis* asserts that the stationary vertex is poorly coupled to the contractile machinery, so that active force is not efficiently transmitted to it, whereas the remodeling vertex is efficiently coupled to the contractile machinery, analogous to the molecular clutch that couples actomyosin to adhesions in migrating or ingressing cells^66–68^. These hypotheses make opposite predictions about protein localization. The *friction hypothesis* predicts that adhesive, anchoring machinery should be enriched at the stationary vertex, where it would generate the required resistance. The *clutch hypothesis* instead predicts that the adaptor proteins coupling actomyosin to junctions should be enriched at the remodeling vertex, where force is being transmitted.

We first addressed the source of the driving force. Live imaging of tricellular zipping using probes for Myosin II and E-cadherin showed that Myosin II was significantly enriched at the remodeling vertex, particularly during the early stages of zipping **(Fig. 3B, C, D).** We further observed local peaks in Myosin II intensity at the remodeling vertex as the *de novo* BCJ elongated **(Fig. 3E).** Calculating the slope of *de novo* BCJ elongation during the 2 min before and 2 min after each Myosin II peak, we found that the post-peak slope was significantly higher than the pre-peak slope **(Fig. 3F and S2A).** Enrichment of contractile machinery at the remodeling vertex, together with the association between local Myosin II peaks and accelerated zipping, establishes a tension asymmetry that generates a net force displacing the remodeling vertex.

We then turned to the immobility of the stationary vertex and tested the *friction hypothesis*, which predicts enrichment of adhesive machinery at the stationary vertex. Contrary to this prediction, E-cadherin intensity was higher at the stationary vertex in only about half of zipping events, with the remainder showing enrichment at the remodeling vertex instead **(Fig. 3B, C, D).** The absolute level of E-cadherin, therefore, does not determine which vertex remains stationary. We also examined the ratio of Myosin II to E-cadherin intensity at both vertices and observed peaks at the remodeling vertex that were absent at the stationary vertex **(Fig. 3G),** with the post-peak elongation slope again exceeding the pre-peak slope **(Fig. S2B, C).** Given the comparable result obtained from Myosin II peaks alone **(Fig. 3F)** and the inconsistent E-cadherin asymmetry **(Fig. 3D),** we conclude that this effect is driven primarily by Myosin II, and we interpret the Myosin II to E-cadherin ratio as a normalized measure of contractility rather than as evidence for a drag term.

If the stationary vertex is not held in place by drag, an alternative explanation is that the forces acting on this vertex genuinely cancel, so that it sits close to mechanical equilibrium and experiences no net force. We tested this directly using vertex geometry. At a vertex where three junctions meet, mechanical equilibrium requires the three tension vectors to sum to zero, which by Lami’s theorem^69^ implies that T/sin(α) is equal for all three junctions, where T is the tension in a junction and α is the angle opposite to it. Using mean Myosin II intensity along each junction as a proxy for tension, we measured T/sin(α) for every junction meeting at the stationary vertex and remodeling vertex **(Fig. S2D).** We quantified departure from balance in two ways: the coefficient of variation (CV) of T/sin(α) across the junctions at a vertex, and the vector imbalance, defined as the magnitude of the summed tension vectors expressed as a fraction of the total tension (see supplementary information). Both measures are zero for a perfectly balanced vertex. Across zipping events, the median CV was ∼12% and the median vector imbalance was 0.14 at the stationary vertex, whereas the median CV was ∼42% and the median vector imbalance was 0.3 at the remodeling vertex (**Fig. S2E, F).** Because Myosin II intensity is an indirect and noisy proxy for junctional tension, neither measure is expected to reach zero even at a perfectly balanced vertex. For a majority of stationary vertices, the junctional tensions are close to balanced, indicating that this vertex lies near mechanical equilibrium. Its lack of displacement can be accounted for within the force-balance framework without invoking a drag asymmetry.

We next examined the tricellular tight junction proteins Angulin-1 and Tricellulin, which mark the vertex itself. Both proteins accumulated at and moved with the remodeling vertex during zipping **(Fig. 3H, S2G, and Video S4).** To probe the dynamics of this accumulation, we performed fluorescence recovery after photobleaching (FRAP) of Angulin-1 at both vertices^70, 71^ **(Fig. 3I, S2H, and Video S4).** Note that the remodeling vertex does not move appreciably over the 250 sec duration of the FRAP experiment. Surprisingly, the mobile fraction of Angulin-1 was significantly higher at the stationary vertex than at the remodeling vertex, 0.7 ± 0.06 vs. 0.4 ± 0.05, respectively **(Fig. 3J),** while the t_1/2_ did not differ significantly **(Fig. S2I).** Counterintuitively, therefore, the stationary vertex is the more molecularly dynamic of the two vertices, whereas the remodeling vertex is more molecularly stable. This data suggests that the static vertex may be less stably connected to the actomyosin cytoskeleton.

To assess the *clutch hypothesis*, we examined the TJ adaptor protein ZO-1, which links TJ transmembrane proteins to the actomyosin cytoskeleton. Live imaging revealed higher ZO-1 intensity at the remodeling vertex than at the stationary vertex during zipping **(Fig. 3K and Video S5),** and mosaic labeling showed ZO-1 accumulating at the zipping front from the receding cell **(Fig. S2J).** Quantification confirmed that ZO-1 was consistently higher at the remodeling vertex **(Fig. 3L and S2K).** This is the localization predicted by force transmission and the opposite of that predicted by the *friction hypothesis*: an actomyosin-to-junction adaptor is enriched at the vertex that moves, consistent with ZO-1 contributing to the coupling through which contractile force reaches the junction. Consistent with this idea, the product of Myosin II and ZO-1 intensities at the remodeling vertex – a combined measure of local contractility and coupling – showed peaks that preceded increases in the rate of *de novo* BCJ elongation **(Fig. S2L, M).** We note, however, that junction elongation rate did not correlate with the instantaneous magnitude of this product; rather, it was local increases in the signal that were associated with accelerated elongation, suggesting that changes in contractile engagement, rather than its absolute level, accompany vertex displacement.

Taken together, these data suggest that tricellular zipping might be driven by tension asymmetry. Contractile actomyosin is enriched at the remodeling vertex, generating the net force that displaces it, and local increases in contractility there are associated with accelerated elongation of the *de novo* BCJ. The stationary vertex, in contrast, sits close to mechanical equilibrium, and its immobility can be explained by the force-balance framework. Additionally, the molecular composition of the two vertices is consistent with differential force transmission. The remodeling vertex is enriched in ZO-1 and contains a comparatively stable Angulin-1 population, whereas the stationary vertex has lower ZO-1 and a more dynamic, rapidly exchanging Angulin-1 population, thus preferentially coupling the contractile machinery in the receding cell to the remodeling vertex.

### Localized actomyosin dynamics drive tricellular zipping

Next, we asked whether Myosin II asymmetry is functionally important for tricellular zipping. Using super-resolution microscopy, we observed that in the early stages of tricellular zipping, Myosin II symmetry breaking occurred, with higher Myosin II intensity at the remodeling vertex than at the stationary vertex (**Fig. 4A and Video S6**). Spatially, Myosin II maintained a horseshoe-shaped accumulation at the zipping front and moved together with the remodeling vertex. The end of tricellular zipping was marked by a reduction in Myosin II intensity at the remodeling vertex (**Fig. 4A and Video S6**). To investigate whether Myosin II activity in the receding cell is important for efficient tricellular zipping, we perturbed Rho-associated protein kinase (ROCK), which phosphorylates and activates Myosin II, by mosaically overexpressing a dominant negative form of ROCK, 3xGFP-ROCK2 K106A (ROCK DN) (**Fig. 4B**). Interestingly, cells expressing ROCK DN exhibited stellate morphology, often with multiple long extensions, compared with control cells in the same embryo (**Fig. 4C**). As a result, ROCK DN-expressing cells were also significantly less circular (median: 0.56) than control cells (median: 0.67) (**Fig. 4D**). We hypothesized that these long extensions resulted from failed or inefficient zipping, so we performed long-term imaging and characterized zipping defects observed in ROCK DN-expressing cells. In control cells, zipping began with an acute angle of ∼10° at the remodeling vertex. As zipping progressed, the remodeling vertex angle increased continuously, with periods of elevated slopes, and the angle became more obtuse by the end of zipping (**Fig. S3A**). Interestingly, in most ROCK DN-expressing cells (67%, 10/15) the extensions failed to resolve even after ∼50 min of imaging (**Fig. 4E and S3A**), and the vertex angle did not change significantly from the beginning of imaging. In a smaller population of cell extensions (33%, 5/15), changes in vertex angle were observed; however, they were generally mediated either by forces generated by neighboring cells, such as cell division (13%, 2/15), or by loss of asymmetric vertex motion, with both vertices moving simultaneously, 13% (2/15), while in one case, zipping completed (**Fig. 4E and S3A**). These data suggest that Myosin II activity is important for efficient tricellular zipping.

**Figure 4.**
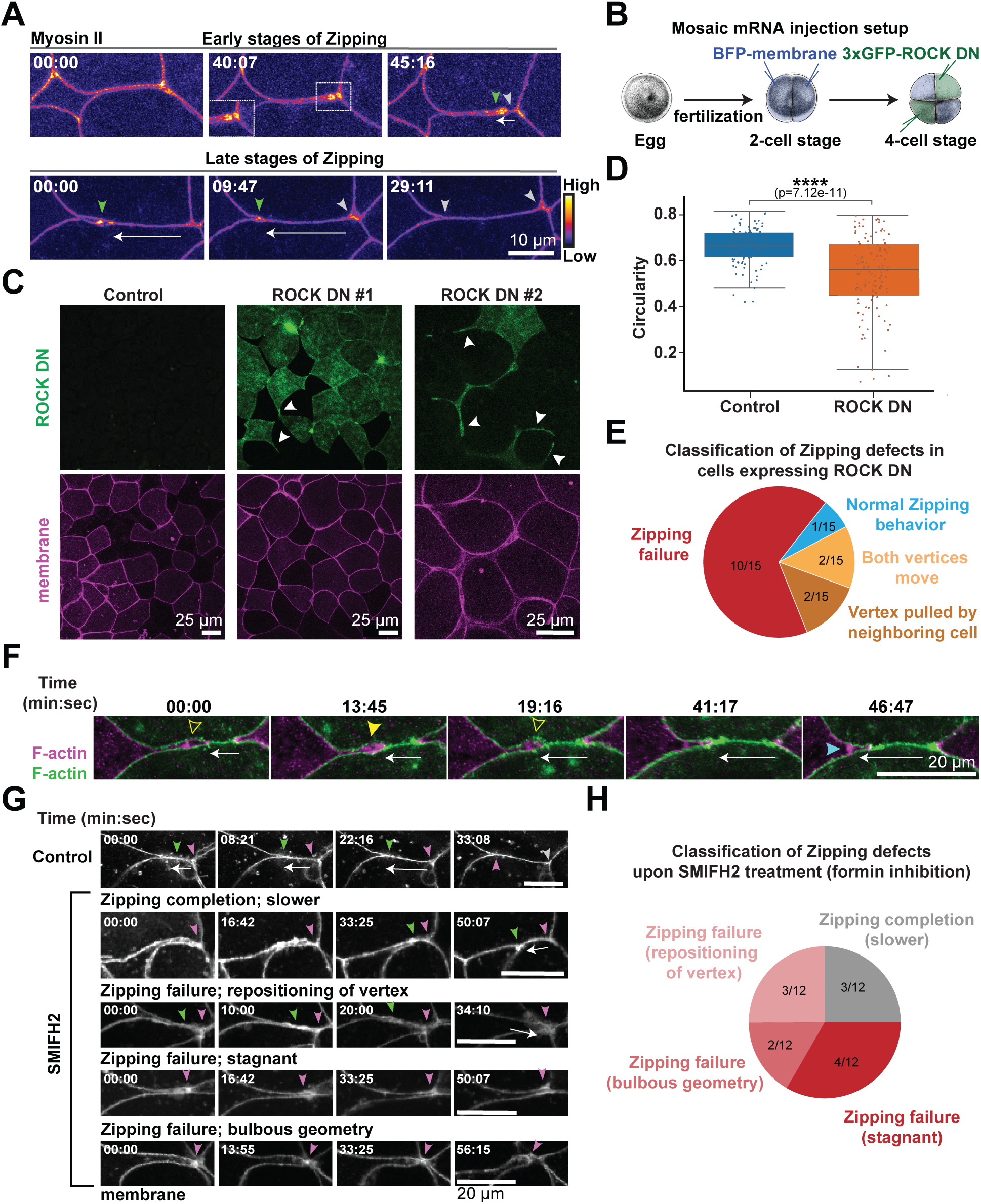
Localized actomyosin dynamics drive tricellular zipping. **(A)** Time-lapse super-resolution confocal images of tricellular zipping in *Xenopus* embryo expressing a probe for Myosin II (SF9-mCherry, FIRE LUT). Horseshoe-shaped accumulation of Myosin II (see inset) moves with the remodeling vertex (green arrowheads). Stationary vertex (gray arrowheads), direction of zipping (white arrows). Time shown in min:sec. Scale bars, 10 µm. **(B)** Experimental setup for mosaic expression of ROCK dominant negative (ROCK DN; 3xGFP-ROCK2 K106A). **(C)** Representative examples of embryos expressing a membrane probe (mem-BFP) without (control) or with ROCK DN (two examples shown). White arrowheads point out unresolved long extensions in cells expressing ROCK DN. Scale bars, 25 µm. **(D)** Quantification of circularity of cells expressing ROCK DN or control cells. Box plots show the median (solid line), the box shows 25^th^ to 75^th^ percentile, and the whiskers represent the min and max values. *n* = 247 control cells and 194 ROCK DN cells, 16 embryos, 3 experiments. Statistical significance was determined using a Mann-Whitney U test; *\*\*\*\*p ≤* 0.0001. **(E)** Pie chart showing the number of cells expressing ROCK DN that exhibited different types of zipping defects. (See Extended Data Fig. S3). **(F)** Time-lapse super-resolution confocal images of tricellular zipping in *Xenopus* embryo mosaically expressing two probes for F-actin (LifeAct-GFP (green) and LifeAct-RFP (magenta)). F-actin accumulates in the receding cell extension in a pulsatile manner (open and closed yellow arrowheads). White arrow indicates the direction of zipping. Time shown in min:sec. Scale bar, 20 µm. **(G)** Representative examples of different kinds of zipping defects observed in embryos treated with 100 µM SMIFH2 compared with 0.2% DMSO (vehicle control). Remodeling vertex (green arrowheads), stationary vertex (magenta arrowheads), direction of zipping (white arrows). Time shown in min:sec. Scale bars, 20 µm.**(H)** Pie chart showing the number of embryos treated with SMIFH2 that exhibited different types of zipping defects (shown in G).

F-actin exhibited distinct dynamics compared with Myosin II. To image F-actin dynamics, we generated embryos mosaically expressing two differently colored fluorescently-tagged F-actin probes. Super-resolution imaging revealed pulsatile F-actin dynamics within the receding cell extension during tricellular zipping (**Fig. 4F and Video S7**). We hypothesize that the pulsatile F-actin accumulation may help pull the zipping membranes together, while additional F-actin accumulation from the zipping cells at the *de novo* BCJ and newly positioned tricellular vertex may stabilize and reinforce the position of the tricellular vertex.

To test whether formin-mediated F-actin polymerization is required for tricellular zipping, we used the formin inhibitor SMIFH2. F-actin intensity at the zipping site differed between SMIFH2-treated embryos and controls; specifically, F-actin signal was either lower than in controls or exhibited higher, stable intensity at junctions (**Fig. S3B, B’**). We hypothesized that these altered F-actin dynamics would affect efficient tricellular zipping. We classified the zipping defects into two main categories. In 25% of cases (3/12) the remodeling vertex moved unidirectionally and a *de novo* bicellular interface formed, but the process took longer than in controls, lasting >50 min, which we classified as slower zipping completion (**Fig. 4G, H**). In the remaining 75% of cases, the long, thin extension failed to resolve and did not generate a *de nov*o BCJ between the zipping cells (**Fig. 4G, H**). In 33% of cases (4/12) the extension remained stagnant for >50 min. In 25% (3/12), movement of the remodeling vertex was observed, but during zipping, the remodeling vertex collapsed with the stationary vertex, resulting in failed zipping. In 16% (2/12), the geometry of the multicellular vertex at the end of the extension changed from the typical acute angle to a bulbous shape. Thus, formin-mediated actin polymerization is important for efficient and successful tricellular zipping. Of note, we did not observe significant accumulation of the Arp2/3 complex (Arp3-mNeon) in the long cellular extension or the zipping cells during tricellular zipping (**Fig. S3C**). Taken together, these results show that localized actomyosin dynamics drive tricellular zipping, and that blocking ROCK or formin activity leads to failed zipping events, resulting in unresolved cell extensions in the developing epithelium.

### Calcium and F-actin pulses are associated with accelerated elongation of the *de novo* bicellular junction during tricellular zipping

To investigate the upstream regulatory pathway that drives pulsatile actomyosin dynamics at sites of tricellular zipping, we examined calcium signaling as fluctuations in intracellular calcium have been shown to promote actomyosin contractility^72–75^. To characterize spatiotemporal dynamics of cytoplasmic calcium during tricellular zipping, we expressed the genetically encoded calcium indicator GCaMP6m^76^. We observed that calcium flashes occurred repeatedly and sporadically within the long cellular extension of the receding cell during zipping **(Fig. 5A and Video S8)**. Although these flashes did not exhibit an obvious periodicity, multiple calcium flashes were consistently observed during tricellular zipping events **(Fig. 5B)**, suggesting a potential functional role for calcium in this process.

**Figure 5.**
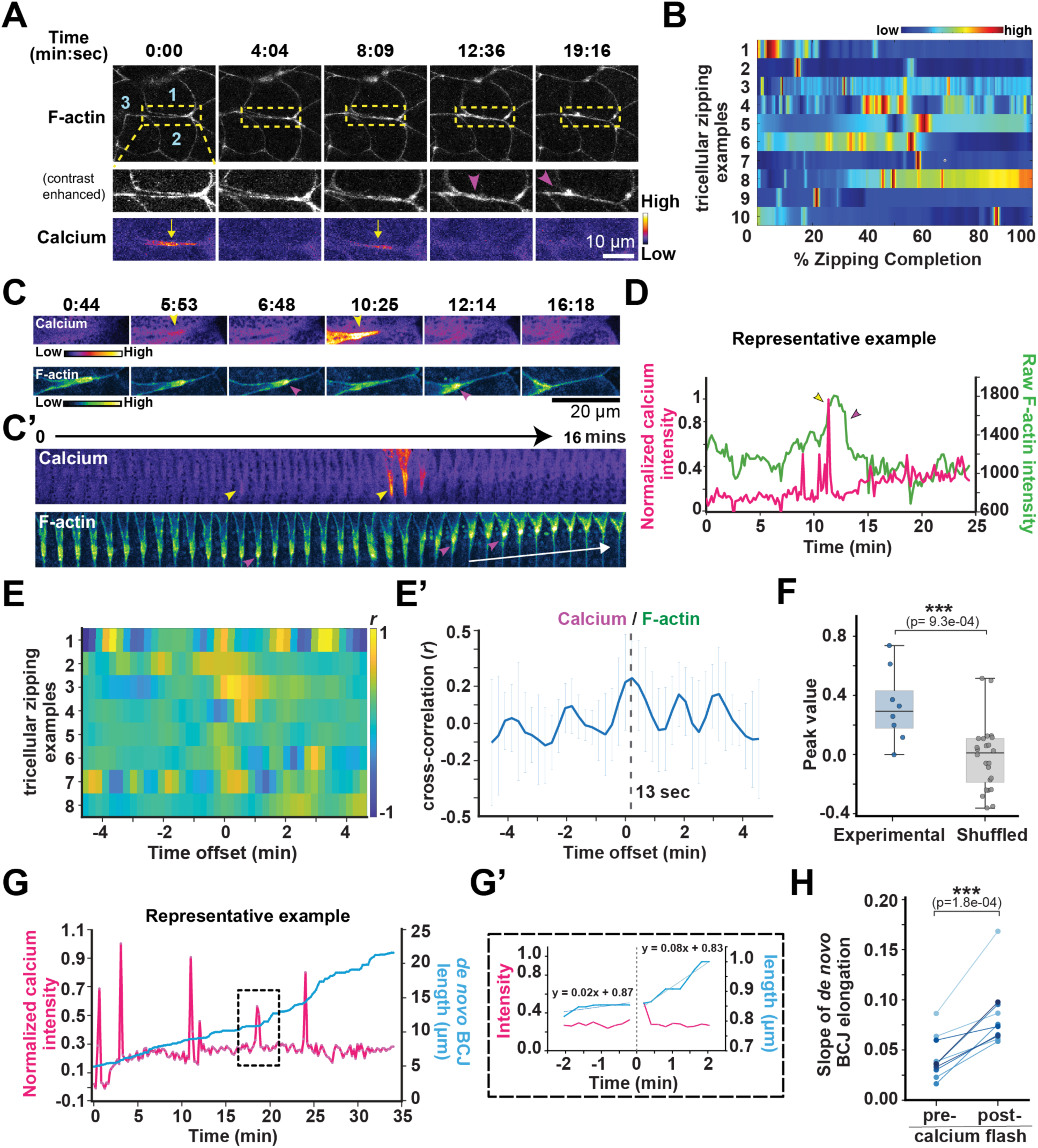
Calcium and F-actin pulses are associated with increased elongation of the *de novo* bicellular junction during tricellular zipping. **(A)** Time-lapse confocal images of a *Xenopus laevis* embryo expressing probes for F-actin, (LifeAct-miRFP703, grayscale) and cytoplasmic calcium (GCaMP6m, FIRE LUT). Zipping cells (cells 1 and 2), receding cell with long extension (cell 3). Enlarged regions (yellow dashed boxes) show repeated increases in cytoplasmic calcium within the long cellular extension (yellow arrows) and F-actin dynamics at the zipping front (magenta arrowheads). Time shown in min:sec. Scale bar, 10 µm. **(B)** Repeated calcium flashes are associated with tricellular zipping across six independent zipping examples. Warm colors indicate increased cytoplasmic calcium intensity. **(C)** Montage of tricellular zipping in a *Xenopus laevis* embryo expressing probes for cytoplasmic calcium (GCaMP6m, FIRE LUT), and F-actin (LifeAct-miRFP703, Green Fire Blue LUT). Scale bar, 20 µm. **(C′)** Kymograph generated from the example shown in (C). Yellow arrowheads indicate localized calcium flashes, and magenta arrowheads indicate corresponding increases in F-actin at the zipping front. The white arrow indicates a sharp elongation of the *de novo* BCJ following a calcium flash. **(D)** Representative plot showing normalized calcium intensity and raw F-actin intensity within the long cellular extension during a single zipping event. **(E)** Heatmap showing the cross-correlation between the rate of change in calcium intensity and the rate of change in F-actin intensity for eight zipping events. The color scale represents the strength and directionality of cross-correlation (positive, negative) across different time offsets. **(E′)** Mean cross-correlation across eight tricellular zipping events, with an average correlation coefficient of *r* = 0.244 and an average offset of 13 s. *n* = 8 zipping events, 7 embryos, 5 experiments. **(F)** Boxplot showing the distribution of *r* values at a 13 s offset for the experimental data (where calcium and F-actin traces are matched) and for shuffled data (where the actin traces were shuffled and paired with different calcium traces as a control). Statistical significance was determined using an unpaired Student’s *t-*test; \*\*\**p* < 0.001. **(G)** Representative plot showing normalized calcium intensity within the long cellular extension, and *de novo* junction length during a single zipping event. (G’) The enlargement shows an isolated calcium flash (boxed region in G) centered at t = 0, with the slope of junction elongation calculated 2 min before and 2 min after the calcium flash. **(H)** Quantification of paired pre-flash and post-flash junction elongation slopes from multiple zipping events. *n* = 10 flashes, 7 zipping events, 7 embryos. Statistical significance was determined using a paired *t*-test; \*\*\**p* < 0.001.

We next asked whether calcium flashes are coupled to F-actin pulses during tricellular zipping. Live imaging showed that calcium flashes were often followed by increases in F-actin intensity within the cell extension during tricellular zipping **(Fig. 5C, C’, D)**. To better quantify this relationship, we measured rates of change in calcium and F-actin intensities within the cell extension. We then performed cross-correlation analysis on these rate-based values **(Fig. 5E, E’)**. Heatmap representations of cross-correlations from individual zipping events revealed variability in cross-correlation patterns, with high *r* value peaks (shown in yellow) offset just to the right of 0 in most samples (**Fig. 5E**). When averaged, there was a cross-correlation between calcium and F-actin, with an average *r* value of 0.244 and an average offset of 13 s. This indicates that increases in calcium signal precede increases in F-actin signal by ∼13 s (**Fig. 5E’**). Statistical analysis confirmed this relationship, as *peak* values at the 13 s offset were significantly higher in the matched calcium/F-actin data than in data where calcium/F-actin pairings were shuffled (**Fig. 5F**). Furthermore, the increase in F-actin level following a calcium flash was restricted to the cell extension terminating at the remodeling vertex. The remodeling vertex exhibited a significant increase in F-actin intensity after a calcium flash, whereas F-actin remained nearly unchanged at the stationary vertex (**Fig. S4A, B**). This observation further supports our force-balance framework (**Fig. 3**), in which the stationary vertex is close to equilibrium and its lack of displacement reflects the absence of active force. Together, these results support a model in which calcium signaling acts upstream of F-actin pulsatile accumulation at the zipping front during tricellular zipping, while the stationary vertex receives no active force, resulting in asymmetric vertex displacement.

Notably, calcium flashes often coincided with elongation of the *de novo* BCJ, raising the possibility that calcium signaling contributes to the progression of tricellular zipping. We quantified calcium intensity together with the length of the *de novo* BCJ during tricellular zipping. These measurements revealed multiple calcium flashes throughout the zipping process, while the *de novo* BCJ progressively increased in length (**Fig. 5G**). Notably, elongation of the *de novo* BCJ often appeared to accelerate after a calcium flash (**Fig. 5G**). To determine whether calcium flashes influence the rate of junction elongation, we quantified isolated calcium flashes, calculating the slope of *de novo* BCJ elongation before and after the flash (**Fig. 5G’**). Across multiple samples, the post-flash slope was significantly higher than the pre-flash slope (**Fig. 5H**). Together, these results show that localized calcium flashes and actomyosin pulses are associated with accelerated tricellular zipping.

### Photochemically induced local calcium increase drives acceleration of tricellular zipping

Having found that calcium flashes are associated with accelerated zipping, we wanted to more directly test whether tricellular zipping is dependent on calcium-mediated actomyosin contractility. We photochemically triggered a calcium burst in the receding cell by calcium uncaging **(Fig. 6A-B, S4C-D, and Video S9)**^77^. The induced calcium bursts were confined to the cell extension; however, their duration and intensity varied. Some receding cells exhibited a more robust, wider, longer-lasting increase in calcium **(Fig. 6A)**, whereas others showed a more localized and shorter-lasting calcium increase **(Fig. S4D)**. We further analyzed F-actin intensity at the site of zipping before and after induced calcium increase. The F-actin intensity increased drastically at the zipping front immediately following the induced calcium burst **(Fig. 6A-D, S4D, and Video S9)**. The magnitude of the increase in F-actin intensity loosely correlated with the duration of the induced calcium burst **(Fig. S4E)**, suggesting that the extent of actomyosin contractility is tunable and depends on upstream calcium signaling during tricellular zipping. Further, we examined the potential molecular clutch, ZO-1, and found that its intensity also increased at the remodeling vertex right after the calcium burst **(Fig. 6E, F)**. To determine whether an increase in actomyosin contractility after induced calcium bursts influences the rate of zipping, we calculated the slope of *de novo* BCJ elongation before and after calcium increase. Across multiple samples, the slope was significantly higher following the calcium burst (**Fig. 6G**), indicating that localized cytoplasmic calcium flashes are sufficient to accelerate tricellular zipping. Thus, our data show that localized calcium flashes within the cell extension promote force generation by increasing actomyosin contractility and enhance force transmission at the remodeling vertex by increasing ZO-1 accumulation, which is crucial for efficient, ratchet-like tricellular zipping behavior. Furthermore, the speed of tricellular zipping can be altered by modulating the timing and duration of calcium flashes.

**Figure 6.**
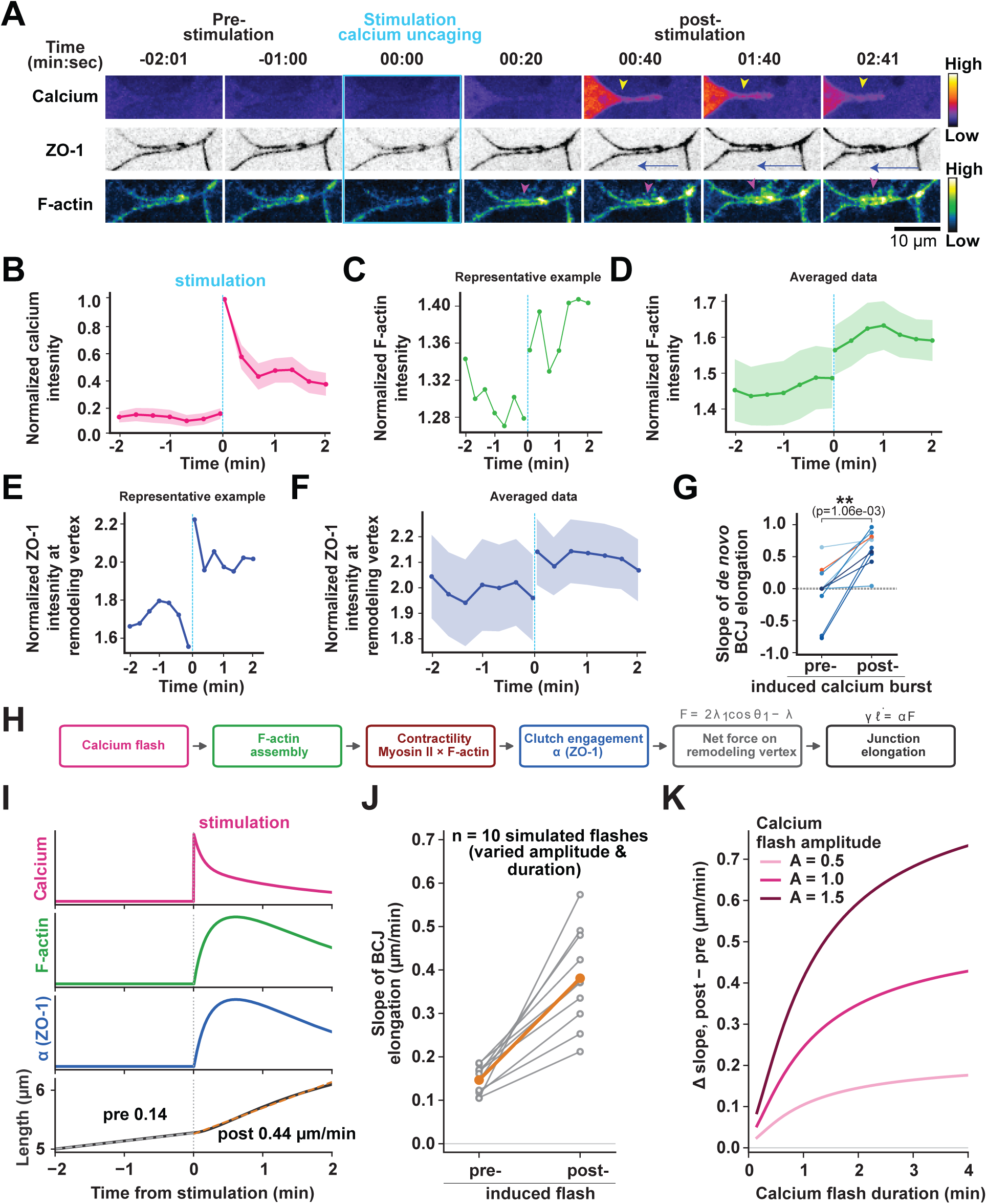
Local calcium increase drives acceleration of tricellular zipping. **(A)** Time-lapse confocal images of tricellular zipping before and after induced calcium bursts in a *Xenopus laevis* embryo expressing probes for calcium (cGAMP6m, FIRE LUT), ZO-1 (mRFP-ZO-1), and F-actin (LifeAct-miRFP703, Green Fire Blue LUT). Time shown in min:sec. Localized stimulation to induce calcium burst was performed at time point 0 (blue box). Yellow arrowheads indicate the increase in calcium, blue arrow indicate the directionality of zipping, and magenta arrowheads indicate increase in F-actin right after the stimulation. Scale bar, 10 µm. **(B)** Normalized calcium intensity at the site of zipping before and after stimulation (blue dotted line). **(C)** Representative plot (quantified from A) showing normalized F-actin intensity (normalized to BCJ) at the zipping site. **(D)** Quantification of normalized F-actin intensity at the site of zipping before and after stimulation. **(E)** Representative plot (quantified from A) showing normalized ZO-1 intensity (normalized to BCJ) at the remodeling vertex. **(F)** Quantification of normalized ZO-1 intensity at the remodeling vertex before and after stimulation. For (B-F) Mean (solid line) ± SEM (shading), *n* = 10 zipping events, 10 embryos, 4 experiments. **(G)** Quantification of paired *de novo* BCJ elongation slopes pre– and post-induced calcium burst. Orange line corresponds to the data in A. *n* = 10 induced calcium bursts, 10 zipping events, 10 embryos, 4 experiments. Statistical significance was determined using a paired *t*-test; \*\**p* < 0.01. **(H)** Calcium-gated model of the response of the remodeling vertex to a calcium flash (see Supplementary Information). A flash drives F-actin assembly with response time τf. Contractility is the product of Myosin II and F-actin concentration. Clutch engagement α = χ s(a) follows contractility through a saturating switch, with χ the coupling strength. The transmitted net force F sets the elongation rate of the BCJs through the relation, γ dℓ/dt = αF. **(I)** Simulated response to a single induced flash at t = 0, modeled on the measured calcium transient in **(B)**. From top: calcium, F-actin, clutch engagement α, and junction length. Dashed lines show the least-squares linear fits to the length dynamics over the 2 min before and the 2 min after the flash, from which we compute the slopes reported in **(J)**. **(J)** Pre– and post-flash elongation slopes for ten simulated flashes (gray, individual flashes; orange, mean). See Supplementary Information for parameter choices. **(K)** Simulated calcium flash-induced increase in elongation rate, as a function of flash duration for three flash amplitudes A (A = 1 is the reference flash in I).

### Calcium-gated model of the response of the remodeling vertex to a calcium flash

The force-balance relation explored in Fig. 3 and S2 treats force transmission efficiency and the friction coefficient as fixed. But our data in Fig. 5 and 6A-G show that contractility is pulsatile. Calcium flashes in the cell extension precede F-actin accumulation by roughly 13 s (**Fig. 5E–F**), Myosin II accumulation is comparatively steady while F-actin pulses (**Fig. 4A, 4F, and 5C–D**), ZO-1 at the remodeling vertex rises after an induced calcium burst (**Fig. 6E–F**), and the rate of BCJ elongation increases after both endogenous and induced calcium increase (**Fig. 5H, 6G**). In our model, we therefore let calcium drive the F-actin component of the contractile machinery and let clutch engagement follow contractility. F-actin activity responds to calcium with a finite relaxation time, combines with a steady local Myosin II pool to generate contractile activity, and promotes force transmission through a junctional clutch engagement coefficient associated with adaptor proteins like ZO-1 **(Fig. 6H and Supplementary Information)**. The resulting transmitted force determines the elongation rate of the *de novo* BCJ **(Fig. 6H).** With the sequential interplay of calcium, F-actin, clutch engagement built into the model **(Fig. 6H, I)**, a transient calcium flash was sufficient to produce a sustained increase in elongation rate comparable to that measured after induced bursts in embryos **(Fig. 6G, J)**. The model further predicted that acceleration depends nonlinearly on both flash strength and duration **(Fig. 6K).** Short or weak calcium flashes produced less acceleration because the resulting contractile activity remained below the threshold, whereas stronger or longer flashes crossed this threshold and resulted in increased elongation. The response is saturated for prolonged flashes, consistent with the weak correlation between F-actin accumulation and induced burst duration **(Fig. S4E)**. Collectively, these findings identify calcium flashes as pulsatile signals that generate pulsatile increases in actomyosin-mediated contractility, thereby producing sustained increases in BCJ elongation.

### Calcium signaling in *Drosophila* is required to restore ectoderm cell shape during mesoderm internalization

Given the importance of calcium flashes to facilitate tissue packing during development in *Xenopus* embryos, we next asked what roles calcium flashes play in developing *Drosophila* embryos. *Drosophila* embryos have been shown to exhibit cell-level calcium flashes^78^, and induced calcium bursts generated by uncaging calcium resulted in apical constriction^77, 79^. However, the role of endogenous calcium flashes in the early *Drosophila* embryo remains poorly understood.

Using *Drosophila* embryos expressing the membrane-bound calcium probe GCaMP8s, we identified cells exhibiting transient changes in cytosolic calcium levels during gastrulation first by eye, and then through segmentation^80^ and computational parameters **(Fig. S5A, B, and Video S10).** We initially observed that various cell shape change events occurred in cells after calcium flashes, but it was unclear if these calcium flashes were associated **(Fig. S5A’)**. Quantification of cell properties within a 2-minute window of a flash event confirmed significant changes in area, both apical constriction and apical expansion **(Fig. S5C, C’)**. Given the similar processes of epithelial cell packing during gastrulation in *Drosophila* and *Xenopus* embryos, we quantified morphological changes using cell circularity and cell shape index as we had in *Xenopus* (Fig. 1B, E). The least-circular cells (initial circularity < 0.7) exhibited a significant increase in circularity 1-minute post-calcium flash **(Fig. S5D).** For all events in which mean circularity increased, cell shape index decreased **(Fig. S5E, E’)**, consistent with the cells changing from more elongated and irregular shapes to a more regularly packed morphology during convergent extension in gastrulation.

The effects of transient calcium level changes on cell shape and behavior **(Fig. S5A’)** prompted us to investigate whether manipulating intracellular calcium levels would affect how cells regulate their shape and thereby influence embryogenesis. We first perturbed calcium through pharmacological drug injection to modulate calcium levels uniformly across the embryo. Embryos injected with DMSO (vehicle control) displayed more calcium flashes than non-injected GCaMP control embryos, consistent with previous observations on the effect of physical perturbation on calcium spiking activity^78^. Injection of the intracellular calcium chelator BAPTA-AM reduced the number of these calcium flashes in GCaMP embryos (**Fig. 7A, B**). The reduction of calcium flashes with BAPTA-AM suggests that BAPTA-AM is decreasing available free cytosolic calcium, but morphogenesis proceeded, likely because there was only a moderate impact on calcium dynamics. Injection of the calcium ionophore, ionomycin, elevated calcium levels uniformly across the embryo (**Fig. 7C**). Ionomycin injection at early stage 5 prevented mesoderm invagination initiation and calcium flashes in the ectoderm, whereas injection at stage 6 resulted in failed mesoderm internalization and retained ectodermal calcium flashes ( **Video S11**). These findings suggest that the timing of calcium perturbation plays a critical role in determining the downstream cellular response. Together, the developmental defects observed in ionomycin injected embryos indicate that calcium dynamics are necessary for proper tissue morphogenesis in the early *Drosophila* embryo.

**Figure 7.**
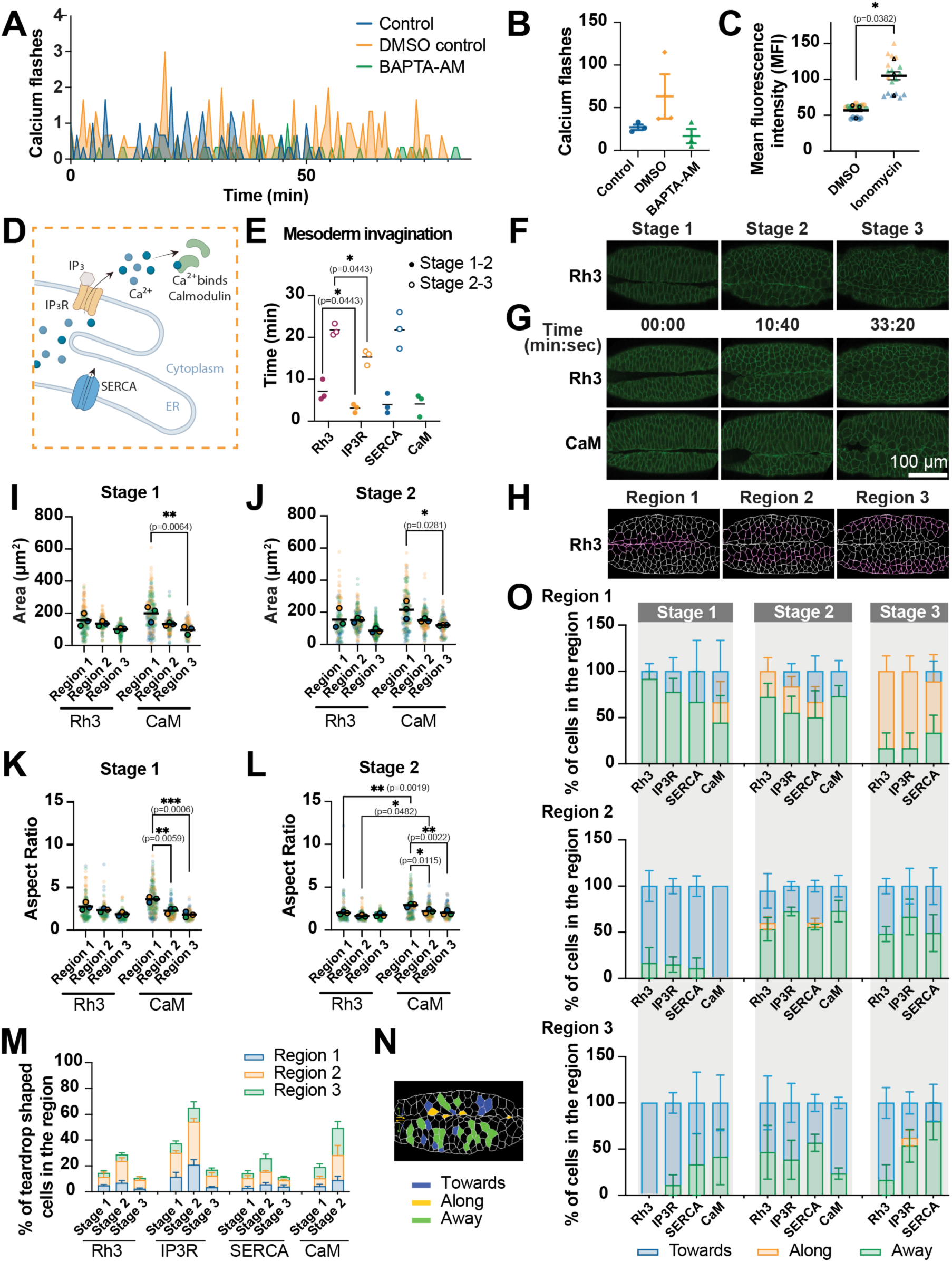
Calcium signaling in *Drosophila* is required to restore ectoderm cell shape during mesoderm internalization. **(A)** Quantification of calcium flash frequency over time. Time point 0 denotes the first recorded calcium flash. Embryos injected with the calcium chelator BAPTA-AM (10 mM) were compared to embryos injected with DMSO (vehicle control) and uninjected control embryos. N = 3 embryos per condition. **(B)** Quantification of total calcium flash number per embryo across treatment conditions. N = 3 embryos per condition. **(C)** Mean fluorescence intensity quantified from 5 regions of interest of equal area per embryo. Embryos injected with the calcium ionophore Ionomycin (1 mM) were compared to embryos injected with DMSO (vehicle control). Colors represent independent biological replicates. N = 3 embryos per condition. Statistical significance was determined using an unpaired t test; \**p* ≤ 0.05. **(D)** Schematic of the intracellular calcium signaling pathway in *Drosophila* embryo. **(E)** Duration of stage transitions (stages 1–2 and stages 2–3) in each genotype. N = 3 embryos per genotype. Statistical significance was determined using a Tukey’s multiple comparison test; \**p* ≤ 0.05. **(F)** Representative images from Rhodopsin (Rh3) RNAi defining ventral furrow stages used for quantitative analyses in panels E, I-M, and O. **(G)** Representative images of mesoderm internalization processes in *Drosophila* embryos expressing the membrane marker P4M (green) with Rh3 and Calmodulin (CaM) knocked down by RNAi. **(H)** Representative images from Rhodopsin (Rh3) RNAi illustrating the definition and boundaries of embryonic region categories used for quantitative analyses in panels I-M and O. **(I–J)** Quantification of cell area by embryonic region in control and CaM RNAi knockdown embryos at stage 1 (I) and stage 2 (J). N = 3 embryos per genotype, with a minimum of 23 cells quantified per embryo. Statistical significance was determined using a Tukey’s multiple comparison test; \**p* ≤ 0.05, \*\**p* ≤ 0.01. **(K–L)** Quantification of cell aspect ratio by embryonic region in control and CaM RNAi knockdown embryos at stage 1 (K) and stage 2 (L). N = 3 embryos per genotype, with a minimum of 23 cells quantified per embryo. Statistical significance was determined using a Tukey’s multiple comparison test; \**p* ≤ 0.05, \*\**p* ≤ 0.01, \*\*\**p* ≤ 0.001. **(M)** Frequency of teardrop-shaped cells across embryonic regions and developmental stages. **(N)** Representative images illustrating the definition of the orientations of teardrop cells in relation to the ventral furrow. **(O)** Frequency distribution of teardrop cell orientations (toward, along, or away from the ventral furrow) with different calcium perturbations.

We next sought to alter calcium signaling through knocking down specific calcium regulators via RNAi. We systematically examined the effects of knocking down the Inositol 1,4,5-trisphosphate receptor (IP_3_R), Calmodulin (CaM), and the Sarco/Endoplasmic Reticulum calcium ATPase (SERCA). Briefly, IP_3_R is an IP_3_-gated calcium channel on the ER membrane responsible for releasing calcium ions from the ER lumen into the cytoplasm, CaM binds free cytoplasmic calcium ions to form a complex that activates downstream cellular activities, and SERCA recycles free cytoplasmic calcium ions back into the ER **(Fig. 7D).** Knockdown of IP_3_R resulted in more rapid mesoderm internalization compared with control (Rh3 knockdown) **(Fig. 7E)**. For reference, we defined the stages of mesoderm internalization as follows: stage 1, furrow forming; stage 2, furrow touching on both sides; stage 3, furrow sealing **(Fig. 7F)**. In contrast to IP_3_R knockdown, CaM knockdown, which would likely more broadly arrest downstream calcium signaling, resulted in severe defects in ventral furrow formation, with the mesoderm failing to fully seal to stage 3 **(Fig. 7G).** Overall, these data suggest that alterations in calcium signaling modulate gastrulation movements.

Our analysis of calcium flashes in *Xenopus* demonstrated that calcium restores distorted epithelial packing to a more hexagonal array. Therefore, to assess the impact of calcium-related gene perturbations on cell packing during *Drosophila* gastrulation, we 1) analyzed cell shape in different regions relative to the invaginating mesoderm, and 2) analyzed the frequency of ‘teardrop-shaped cells’. We divided segmented embryos into 3 regions relative to the ventral midline: region 1 comprised cells in direct contact with the invaginating cells; region 2 comprised cells adjacent to region 1; and region 3 comprised all remaining cells, with minor adjustments based on total cell area **(Fig. 7H)**. CaM knockdown resulted in significantly greater differences in area and aspect ratio between regions compared to Rh3 knockdown controls **(Fig. 7I-L)**. Furthermore, cells in CaM knockdown regions 1 and 2 displayed significantly larger aspect ratio compared to the corresponding regions in Rh3 knockdown embryos **(Fig. 7L)**, indicating that cells in CaM knockdown regions 1 and 2 are more elongated during stage 2. In IP_3_R knockdown embryos, region 2 showed significantly higher cell area **(Fig. S5F)** and aspect ratio **(Fig. S5G)** compared to other regions during stage 3. Overall, these data suggest that calcium signaling is required to restore cell shape following mesoderm invagination. Both IP_3_R and CaM knockdown embryos displayed an increased frequency of teardrop-shaped cells in regions 2 and 3 during stage 2 **(Fig. 7M).** Each knock-down displayed a distinct directionality profile of teardrop-shaped cells, and an increased proportion of these cells oriented towards the ventral furrow was observed in IP_3_R, SERCA, and CaM knockdown embryos compared to the Rh3 control **(Fig. 7N, O)**. Overall, the analysis of ectodermal cell shape during mesoderm internalization shows that calcium signaling plays a role in promoting hexagonal epithelial packing and the resolution of acute-angled junctions.

## Discussion

Epithelial morphogenesis and homeostatic processes are often facilitated by changes in spatial arrangement of individual cells within the tissue. Tightly regulated crosstalk between actomyosin dynamics and junctional rearrangement promotes changes in cell packing while maintaining barrier function. However, the spatiotemporal regulation of upstream signals that drive local junctional remodeling across different physiological contexts in proliferative epithelial tissues remains unclear. In this study, we uncover a previously uncharacterized process of junction remodeling, which we term ‘tricellular zipping,’ that resolves cell extensions generated by ingression of the cytokinetic furrow and promotes ordered epithelial packing in *Xenopus* embryos (Fig. 1, 2, and S1). Tricellular zipping resolves a long, thin extension terminating at a transient higher-order junction through asymmetric displacement of a tricellular vertex and formation of a *de novo* bicellular interface (Fig. 1, 3). We also observed tricellular zipping in E7.5 mouse embryos (Fig. 1D), suggesting that it may represent a broadly conserved mechanism for facilitating tissue packing in rapidly proliferating tissues. Combining modeling and imaging approaches, we find that asymmetric displacement of the remodeling vertex requires actomyosin force generation and effective force transmission to junctions. In contrast, the stationary vertex remains close to mechanical equilibrium and therefore undergoes no net displacement (Fig. 3, 4, S2, and S3). Furthermore, we discovered that upstream calcium signaling drives localized, pulsatile actomyosin contractility required for efficient zipping (Fig. 5, 6, and S4). Extending our observations beyond *Xenopus*, we show that disrupting calcium signaling in *Drosophila* embryos affects ectodermal cell shape and packing during mesoderm internalization (Fig. 7 and S5). Our combined multidisciplinary and multisystem approach uncovers a new mode of junction remodeling and highlights the importance of transient calcium signaling in driving junctional rearrangement to facilitate morphogenesis.

Early embryogenesis is associated with rapid cell divisions that alter local topology, requiring epithelial cells to rearrange while maintaining barrier function. In the rapidly dividing *Xenopus* blastula epithelium, we show that tricellular zipping resolves the long extensions formed as neighboring cells are pulled in by the cytokinetic furrow (Fig. 2 A, B). The timing and mechanism of tricellular zipping suggest that it is coordinated with, but temporally distinct from, cytokinesis. The basolateral-to-apical progression of tricellular zipping may reflect the geometry imposed by cytokinetic furrow ingression, which also proceeds from basal to apical^19, 65, 81^, and may determine apicobasal differences in cortical actomyosin organization or the sequential maturation of junctional complexes. Thus, tricellular zipping appears to function as a corrective mechanism for cell division-induced packing defects. Determining how cytokinesis establishes the initial cell geometry and molecular asymmetry that subsequently directs zipping will be important areas for future research. Our study also lays the foundation for exploring the role of calcium signaling in driving changes in local packing geometry following cytokinesis in other systems including *Drosophila* epithelial tissues^16–18, 63, 82^.

Our study shows that upon resolution of cell division-induced cellular extensions, the individual cell circularity increases, and cells become more hexagonal (Fig. 1E). At the tissue scale, this process facilitates packing changes during the *Xenopus* blastula-to-gastrula transition, when mean cell circularity increases and the cell shape index decreases (Fig. 1B), suggesting that the developing *Xenopus* epithelium may transition from a fluid-like to a solid-like state, consistent with previous reports in other systems^3, 8, 83–85^. Whether tricellular zipping is restricted to resolving cytokinesis-induced cellular extensions remains unknown. Establishing the frequency of tricellular zipping in other developing tissues and determining whether related behaviors occur in adult homeostatic or regenerating epithelia will broaden our understanding of this process.

A key feature of tricellular zipping is the asymmetric force generation and unidirectional movement of the remodeling vertex. This feature is common with several previously characterized junction remodeling processes including T1 transitions^47, 48, 51^, vertex sliding^50^, and other types of asymmetric vertex displacement^52, 53, 86^, but we show that tricellular zipping is geometrically and mechanically distinct. During a T1 transition, one BCJ contracts into a four-cell vertex, and a new junction forms in the orthogonal direction, while during vertex sliding, one vertex moves along an existing cell-cell contact. In contrast, tricellular zipping initiates in cells with a pre-existing extension, and the *de novo* bicellular interface forms through membrane convergence (Fig. 1 and S1F). Consistent with these distinctions, zipping occurs in cells that exhibit lower circularity and have cell extensions terminating at an acute vertex angle (Fig. 1E). Mechanistically, we show that the immobility of the stationary vertex is determined by a force balance framework and not because of high adhesion-mediated friction (Fig. 3 and S2). This differs from asymmetric vertex movement in Caco-2 cells, in which a force-mediated increase in E-cadherin stabilizes the less motile vertex^53^. Future studies should determine whether local mechanical heterogeneity driven by lateral cis-clustering of E-cadherin contributes to movement of the remodeling vertex, as previously reported during convergent extension in *Xenopus* embryos^52^. Notably, we observed no intracellular calcium flashes during vertex sliding, which contrasted with tricellular zipping (Fig. S4F), suggesting that distinct signaling mechanisms govern different vertex remodeling processes. These observations expand the known mechanisms by which epithelia reorganize their cell-cell junctions. Different vertex remodeling mechanisms may be utilized based on local cell geometry and the origin of packing defects. Future studies should determine how these processes act in concert, how tissues select among them, and whether geometric constraints determine whether a junction undergoes vertex sliding or tricellular zipping.

Previous studies have shown that ZO-1 undergoes phase separation and supports nascent tight junction formation^87–91^. We propose that a similar phase separation mechanism might help assemble the remodeling tricellular vertex, enabling junctional complexes to remain associated as the remodeling vertex moves. Our quantification showed that the elongation rate of the *de novo* BCJ correlated with local increases in the product of ZO-1 and Myosin II signals, rather than an absolute threshold, suggesting that vertex displacement depends on the engagement of the contractile apparatus. Perturbing ZO-1, Angulin-1, or Tricellulin is needed to distinguish their contributions to force transmission from their roles in junction organization and barrier integrity. Other junctional proteins like Cingulin, Paracingulin, Vinculin, and catenins also bind actomyosin at apical junctions^32, 92, 93^, and it would be interesting to investigate whether they may act along with ZO-1 as a molecular clutch.

Although junction remodeling commonly involves crosstalk between actomyosin and junctional proteins, the upstream signaling mechanisms directing localized force generation remain unclear. We show that localized, transient calcium flashes within cellular extensions drive pulsatile F-actin accumulation and accelerate BCJ elongation during tricellular zipping (Fig. 5). Calcium flashes were localized to the cell extensions and selectively increased F-actin at the remodeling vertex, and not at the stationary vertex (Fig. S4A, B), explaining how transient signals produce persistent asymmetric vertex movement. Using photochemically induced calcium increase, we show that inducing calcium bursts increased actomyosin contractility and ZO-1 accumulation at the remodeling vertex and immediately accelerated zipping (Fig. 6A–G). The pulsatile calcium and F-actin dynamics support a ratchet-like model in which calcium triggers cycles of actin assembly and contractility, potentially through Rho signaling^75, 94, 95^ or calcium-responsive actin modulators such as the formin INF2^96, 97^. Consistently, dominant negative ROCK caused unresolved stellate extensions and impaired zipping, while formin inhibition slowed or stalled zipping (Fig. 4), indicating that ROCK-dependent contractility and formin-dependent actin polymerization are required. An important unresolved question is how calcium is periodically and locally elevated within these long cellular extensions. Increased membrane tension within the extensions may activate mechanosensitive calcium channels^98, 99^. Alternatively, a high concentration of membrane-associated endoplasmic reticulum within the extensions could produce pulsatile calcium release^100^. Identifying the source of these transient intracellular calcium flashes, and defining how calcium interacts with downstream effectors are important directions for future work.

Our findings provide the first evidence that localized calcium flashes act as upstream spatiotemporal regulators of actomyosin-dependent asymmetric vertex remodeling, linking localized calcium activity to persistent changes in cell shape and packing. Although tricellular zipping facilitates packing transitions, we found it challenging to analyze tissue-scale consequences in *Xenopus* due to frequent embryonic lethality upon calcium perturbation. We therefore asked whether the spatiotemporal regulation of calcium flashes similarly influences epithelial packing in another organism: *Drosophila* embryos. Calcium flashes during early *Drosophila* embryogenesis (2-4 hours after egg laying) have been documented^78^. Furthermore, ectopic calcium release in *Drosophila* epithelia is associated with apical constriction^77^. However, the function of endogenous cytosolic calcium flashes was poorly understood. Using the genetically encoded calcium sensor GCaMP, we visualized calcium flashes coinciding with the onset of gastrulation in *Drosophila*. Notably, we were unable to detect subcellular events associated with zipping; instead, we mainly observed calcium flashes that filled the entire cell. Focusing on the ventral side of the *Drosophila* embryo, as mesoderm internalization is accompanied by continuous actomyosin dynamics and junction remodeling in this region, we found that calcium flashes occurred consistently in ectoderm cells after the onset of apical constriction and internalization of the mesoderm. Flashes correspond to regions of the ectoderm that are stretched by the mesoderm and where EGFR-dependent^101^ and force-dependent^102, 103^ Myosin II recruitment occurs.

Injection of the vehicle control DMSO alone increased the number of calcium flashes. This observation is consistent with previous reports showing that injection of water into *Drosophila* embryos promotes calcium flashing behavior^78^. Interestingly, pricking the embryo without delivering any solution did not alter flash frequency, suggesting that the addition of volume and a mechanical or osmotic perturbation is responsible for the increased calcium flashing. Pharmacological perturbation of calcium dynamics using either the calcium chelator BAPTA-AM or the calcium ionophore ionomycin produced results consistent with our expectations. BAPTA-AM reduced the elevated flashing frequency induced by DMSO injection, whereas ionomycin elevated intracellular calcium levels. Regardless of the timing of injection, ionomycin caused mesoderm internalization failure. These results suggest that forcing baseline calcium levels to remain excessively high impairs mesoderm internalization.

To further investigate the function of calcium on embryonic development, we depleted *Drosophila* embryos of genes encoding key components of calcium signaling pathways. We conclude that calcium signaling in the ectoderm modulates cells’ responses to mesoderm internalization. This conclusion is based on the following: 1) Calcium flashes and increased calcium levels are most prominent in the ectoderm. 2) IP_3_R depletion resulted in more efficient mesoderm internalization, whereas CaM depletion caused mesoderm internalization defects and defects in ectoderm cell shape. 3) Injection of ionomycin after cellularization stalls mesoderm invagination. 4) Similar to what we observed in *Xenopus* embryos, we identified ectoderm cells with acute vertex angles, which we refer to as ‘teardrop-shaped cells’ in the *Drosophila* embryo, although we did not observe the tricellular zipping behavior seen in *Xenopus*, suggesting potential species-specific differences in the cell functional roles. Whether *Drosophila* teardrop-shaped cells represent cellular extensions being pulled by other cells or cellular protrusions generated by the teardrop-shaped cells themselves requires future investigation. The frequency of teardrop-shaped cells was increased in IP_3_R mutant embryos across stages and regions. Overall, each genotype exhibited a distinct pattern of teardrop-shaped cell orientation, suggesting that calcium signaling at different stages of calcium processing may fine-tune tissue dynamics during mesoderm internalization. Further work is needed to address whether teardrop-shaped cells result from mesoderm pulling. Given that our earlier injection experiments demonstrated that microinjection was sufficient to elevate calcium flash frequency in *Drosophila* embryos, mechanical force could be an upstream regulator of calcium signaling.

Taken together, using a multidisciplinary and multisystem approach we show a broadly relevant mechanism by which transient calcium flashes drive junctional remodeling to maintain epithelial packing during morphogenesis. These findings establish a framework for exploring how localized calcium signaling regulates junctional dynamics in other proliferative developmental and homeostatic tissues, with broader implications for functional tissue organization and physiology.

## Methods

### Experimental models

#### Xenopus laevis

Adult *Xenopus laevis* female frogs (wild type and albino) and wild type male frogs were purchased from *Xenopus* 1 (Dexter, MI) or the National *Xenopus* Resource (NXR, Woods Hole, MA). Female frogs were injected with human chorionic gonadotropin (HCG, MP Biomedicals, Irvine, CA) to induce them to lay eggs. Male frogs were used for acquisition of testes for sperm preparations. Frogs were housed in a recirculating tank system (Tecniplast, Milan, Italy), which constantly monitors water quality parameters (temperature, pH, and conductivity) to ensure consistent water quality for an optimal environment for frog health. Daily health and maintenance checks were performed by Animal Care Staff.

All studies strictly adhered to the compliance standards of the US Department of Health and Human Services Guide for the Care and Use of Laboratory Animals and were approved by the University of Michigan’s Institutional Animal Care and Use Committee. A board-certified Laboratory Animal Veterinarian oversees our animal facility.

#### Drosophila melanogaster

The following stocks of *Drosophila melanogaster* were used in the study: *Drosophila melanogaster w; matGAL4-VP16 –67; matGAL4-VP16 –15* (Martin Lab stock collection), *Drosophila melanogaster UAS-myr-GCaMP8s* (Gift from Dr. Troy Littleton, MIT), *Drosophila melanogaster w; mat67; mat15, gap43::mCherry7/TM3, Ser*^104^, *D. melanogaster:* 67SqhGFP/Cyo;15UASpmCherry-P4M/TM3^105^, *D. melanogaster:* GCaMP8s. attP2 8622 y1 w67c23; P{CaryP}attP2 68A4 (Littleton lab stock collection), *D. melanogaster*: y[1] sc[*] v[1] sev[21]; P{y[+t7.7] v[+t1.8]=TRiP.GL01052}attP2 (36885), *D. melanogaster*: y[1] sc[*] v[1] sev[21]; P{y[+t7.7] v[+t1.8]=TRiP.HMS01318}attP2 (34609), *D. melanogaster*: y[1] v[1]; P{y[+t7.7] v[+t1.8]=TRiP.GLC01786}attP40 (51686), *D. melanogaster*: y[1] sc[*] v[1] sev[21]; P{y[+t7.7] v[+t1.8]=TRiP.HMS02878}attP2 (44581) (Bloomington *Drosophila* stock center).

The flies were housed in plastic vials containing standard cornmeal medium and kept at room temperature (∼22°C) in a humidified fly room (65% humidity). Flies were transferred to fresh vials on a weekly basis to ensure optimal conditions. The flies used in experiments were kept in a dedicated incubator at 25°C with 65% humidity, in mating cages placed on apple juice agar plates supplemented with yeast paste.

### Mice

All mouse experiments were performed in accordance with University of Michigan Institutional Animal Care and Use Committee requirements and Unit for Laboratory Animal Medicine guidelines. The day a vaginal plug was observed was designated embryonic day (E)0.5. Mice were homozygous for the *Tjp1^tm1Tlch^*allele (ZO-1–GFP^106^) and maintained on a mixed genetic background.

### Embryo handling and microinjection

#### *Xenopus laevis* embryos and microinjection

After egg collection, *Xenopus* eggs were fertilized *in vitro*, dejellied in 2% cysteine in 1X Mark’s Modified Ringer’s solution (MMR) (5 mM HEPES, 100 mM NaCl, 2 mM KCl, 1 mM MgCl_2_, 2 mM CaCl_2_, pH 7.4), pH 7.8 and stored in 0.1X MMR (Woolner et al., 2009). A 5 nl volume containing the desired mRNAs was injected into the animal hemisphere of the embryos four times at either the 2-cell or 4-cell stage. Embryos were kept in 0.1X MMR overnight at 15°C or 13°C and then imaged starting at blastula stage (Nieuwkoop and Faber stage 9)^64^ by live microscopy techniques. Each 5 nl injection contained the following amount of mRNA: mem-BFP: 100 pg for early-stage imaging, mem-BFP: 60 pg^107^; mCherry-farnesyl: 50 pg^108^; LifeAct-miRFP703: 150 pg^109^; MgcRacGAP-3xGFP: 2 pg^110^; SF9-mCherry: 74 pg^21^; E-cadherin-3xGFP: 25 pg^19^; TagBFP2-ZO-1: 60 pg^111^; mRFP-ZO-1: 200 pg^19^; Angulin-1-3xGFP: 25 pg^19^; GFP-Tricellulin: 2 pg^19^; LifeAct-RFP: 25 pg^21^; LifeAct-GFP: 25 pg^19^; Arp3-mNeon: 35pg (this paper); GCaMP6m: 500 pg^75^. For mosaic expression of ROCK DN, cells were injected twice per cell when embryos were at 2 –cell stage, with 60 pg mem-BFP. The embryos were allowed to develop to 4-cell stage at room temperature. Two cells located diagonally from each other were injected with 30 pg 3xGFP-ROCK2 K106A (ROCK DN) (Gift from Dr. Bill Bement, University of Wisconsin, Madison).

#### *Drosophila melanogaster* embryos and microinjection

To create RNAi knockdowns, female flies carrying maternal tubulin GAL4 mat67 and mat15 transgenes^112^ with both sqh::GFP and mCherry::P4M^105^ were mated with males that carried the target gene RNAi construct^113^. The female offspring were screened to exclude balancer chromosomes and were then crossed with their sibling males. The embryos produced from these crosses were subsequently used for microinjections and live time-lapse imaging. Unless stated otherwise, mating cages were kept at 25°C.

The embryos were dechorionated, mounted using embryo glue, and then dried out for 5 minutes using Drierite before being covered with a 3:1 mixture of halocarbon 700 and 27 oils. Using a microinjection system from World Precision Instruments, the embryos were then injected with either BAPTA/AM (10 mM, 196419-25MG, Sigma-Aldrich, Burlington, MA) or Ionomycin (1 mM, I0634-1MG, Sigma-Aldrich, Burlington, MA), both dissolved in DMSO. A DMSO-only injection was used as the vehicle control for all drug injection experiments.

### mRNA preparation for *Xenopus* microinjections

To transcribe mRNA *in vitro*, all DNA constructs were linearized using NotI-HF or KpnI (for theTagBFP2-ZO-1 construct) except for LifeAct-miRFP703 (which was not linearized), then transcribed with the mMessage mMachine SP6 Transcription Kit (AM1340, Thermo Fisher Scientific, Waltham, MA), and purified with the RNeasy Mini Kit (74104, Qiagen, Germantown, MD, USA). After purification, mRNA size was confirmed by running it on a 1% agarose gel with 1% bleach. mRNA was stored at –80°C until use.

### Microscope image acquisition

#### Xenopus laevis embryo imaging

Embryos were suspended in 0.1X MMR or drug solution in 0.1X MMR and mounted in a custom 0.8 mm-thick metal slide with a 10 mm hole in the center; embryos were sandwiched between two glass coverslips adhered with vacuum grease for imaging.

#### Xenopus long term early-stage imaging

Time lapse images during the blastula-to-gastrula transition (Fig.1A, 2A, B) were captured using an inverted Olympus FluoView 1000 confocal microscope with mFV-10-ASW software, equipped with a UPlan FL N 40XO objective (NA = 1.3, working distance = 0.22 mm). Movies were acquired for mem-BFP by scanning the 50 apical Z-planes (step size of 0.5 μm) of a 512 x 512-pixel area (1X zoom) with a 405 nm laser at 8 μs/pixel every 5 min.

#### Tricellular zipping imaging

Movies were generally acquired using an inverted Olympus FluoView 1000 confocal microscope (Fig. 2C, 3I) with mFV-10-ASW software, equipped with a UPlan FL N 60XOS objective (NA = 1.4, working distance = 0.12 mm) or Olympus FluoView 3000 (Fig. 1 C, 2G, 3B, 4C, 4G, S1D, S1F, S2C, S2F, S3A, S3B, and S3G), equipped with Plan Apo N 60X OSC objective (NA = 1.4, working distance = 0.12 mm), with FV31S-SW version 2.6.1.243 software (Powered by H-PF Version 2.14.2.717). Movies were acquired by sequentially scanning the ∼10 apical Z-planes (step size of 0.5 μm) of a 512 x 512-pixel area with a 405 nm, 488 nm, 568 nm or 647 nm laser at 4 or 8 μs/pixel.

#### FRAP image acquisition

Time-lapse movies were acquired for FRAP experiments by scanning the Z-plane with the brightest signal of a 250 x 250-pixel area (2X zoom) with a 488 nm laser at 8 μs/pixel. Photobleaching was performed using the SIM scanner with the clip tornado function of the Olympus FV1000 mFV-10-ASW software. Videos were acquired by imaging for 5 frames (for a pre-bleach baseline), and then a 30% 405-nm laser was pulsed in a circular region (6 µm in diameter) that encompassed the vertex of interest and the neighboring cytosol for 600 msec^71^.

#### Super-resolution imaging

Super-resolution imaging of tricellular zipping in *Xenopus* embryos (Fig. 4A, F and 5C) was performed on a Zeiss LSM980 confocal microscope equipped with an Airyscan2 detector using a Plan-Apochromat 63x/1.4 oil immersion objective with an optical zoom of 2.0-4.0X. Z-stacks of 5-8 µm were obtained with a 0.16 µm z-steps. The airyscan images were processed using Zeiss Zen blue software (ZEN 3.13). The Wiener-filter for airyscan processing was manually adjusted for each channel to produce the best signal-to-noise ratio images.

#### Calcium imaging

Live calcium imaging videos (Fig. 5) of late blastula-early gastrula-stage *Xenopus* embryos were acquired using the Olympus FluoView 3000 as described above. Embryos were imaged with a scanning speed of 2 μs/pixel, a 512 × 512 field of view, and 2X zoom, capturing 6 of the most apical slices of the animal cap of the embryo, with a step size of 0.5 μm.

#### Drosophila melanogaster imaging

Embryos were dechorionated and positioned ventral-side-up in custom slide chambers. The chambers were built using embryo glue (double-sided tape dissolved in heptane) with #1.5 coverslips (48393-151, VWR, Randor, PA) as spacers and a #1 coverslip (48366-067, VWR, Randor, PA) as covers. Once the embryos were mounted, the chamber was filled with Halocarbon 27 oil. All images were acquired on a Zeiss LSM 900 confocal microscope, with a 40×/1.2 Apochromat water objective at room temperature. Pinhole settings ranged from 1–2 airy units. For two-color, live imaging, simultaneous excitation was used with band-pass filters set at ∼ 488 – 546 nm for EGFP and ∼ 595-700 nm for mCherry. All images shown are representative of at least three embryos unless otherwise stated.

### Mouse embryo preparation and live imaging

Embryos were collected at E7.5 using a dissection procedure adapted from Gredler and Zallen, 2023^114^, with dissections performed in ice-cold phosphate-buffered saline (PBS). Surrounding extraembryonic tissues were removed to expose the intact egg cylinder. Embryos were transferred to prewarmed high-glucose DMEM containing L-glutamine and sodium pyruvate (11995073, Thermo Fisher Scientific) supplemented with 10% fetal bovine serum (25-550, Genesee Scientific, El Cajon, CA). Individual embryos were mounted in a droplet of medium on a glass slide within a SecureSeal imaging spacer (654002, Grace Bio-Labs, Bend, OR). A No. 1.5 coverslip was placed over the spacer to seal the chamber, with the surface endoderm facing the coverslip. Mounted embryos were immediately transferred to a Tokai Hit environmental chamber maintained at 37°C and 5% CO₂.

Live imaging was performed on a Nikon AX R confocal microscope equipped with an NSPARC detector and a 60×/1.42 NA Plan Apo oil-immersion objective. GFP was excited at 488 nm, and emission was collected between 502 and 546 nm. Images were acquired at 1024 × 1024 pixels using resonant bidirectional scanning with 8X line integration. Z-stacks were collected at 0.5-µm intervals every 3 min.

### Dye and drug treatments

To stain the cell membrane, *Xenopus* embryos were incubated in CellMask Deep Red plasma membrane stain (C10046, Thermo Fisher Scientific) at 2 μg/ml in 0.1X MMR for 2 minutes and then washed for 10 minutes in 0.1X MMR before imaging.

SMIFH2 (344092, Millipore Sigma, Burlington, MA) was resuspended in DMSO to a concentration of 10 mM, aliquoted, snap frozen in liquid nitrogen, and stored at –80°C. Immediately before imaging, an aliquot of SMIFH2 was thawed and diluted to 100 μM in 0.1X MMR. Embryos were placed in this solution (or the equivalent amount of DMSO in 0.1X MMR as vehicle control), mounted on a slide, and imaged for up to 1 hour.

### Calcium uncaging

DMNP-EDTA (caged calcium), AM ester (50051, Biotium, Fremont, CA) was resuspended in DMSO to a stock concentration of 50 mM and used on the same day. The stock solution was further diluted to 0.5 mM in 0.1X MMR, and embryos were incubated in this solution for 30 min. All handling of the compound was performed under complete darkness with the assistance of night vision goggles. Following incubation, embryos were washed twice in 0.1X MMR and mounted in an imaging chamber under dark conditions. Time-lapse images of GCaMP6m, mRFP-ZO-1, and LifeAct-miRFP703 (F-actin) were acquired by sequentially scanning 6 apical *z*-planes with a step size of 0.50 μm. Images were acquired from a 512 × 512-pixel field at 1.5X zoom using 488 nm, 546 nm, and 647 nm lasers, with a scanning speed of 2 μs/pixel. Imaging was performed for 5 min using an Olympus FV3000 microscope equipped with aPlan Apo N 60X OSC objective (NA = 1.4, working distance = 0.12 mm). For Ca²⁺ uncaging, a rectangular region of interest encompassing the zipping junctions was created and stimulated with a 405 nm laser at 30% power, a pixel dwell time of 10 μs/pixel, and an exposure time of 1929.30 ms. Following stimulation, time-lapse imaging was continued for an additional 10 min.

### Quantification and statistical analysis

All experiments were conducted using three or more biological replicates. The *n* (number of cells analyzed), N (number of embryos) values and experimental days are shown in each figure legend. Statistical tests were chosen following guidance from the Handbook of Biological Statistics^115^. For all the *Xenopus* data, distribution was tested based on normality and equal variances in python (package: scipy (1.17.0)). Then based on the result, either a Student’s t test, Welch’s t test, or a Mann–Whitney U test was used. Specific test and corresponding ‘*p*’ values are represented in the figure and figure legend. Graphs were plotted in python >3.8 using the following packages: matplotlib (3.10.8), numpy (2.4.2), openpyxl (3.1.5), pandas (2.3.3), pip (25.0.1), scipy (1.17.0), seaborn (0.13.2), statsmodels (0.14.6), and statannotations (0.7.2). For *Drosophila* data, statistical analysis was performed in GraphPad Prism 11.0.0.

#### Xenopus laevis quantification

All *Xenopus* images in the figures were processed (e.g., max projected) across each channel and were manually adjusted to show relevant features in Fiji (ImageJ) 2.16.0/1.5p^116^. Unless otherwise stated, all post-acquisition adjustments are consistent between control and treatment conditions within an experiment. All quantification was performed using Fiji; basic data handling was performed using Microsoft Excel 365; plotting and statistical analysis was performed using python >3.8, R (2025.09.1+401, Open source using ggplot2 and dplyr packages), or MATLAB (The MathWorks Inc.) as specified.

#### Quantification of circularity and cell shape index during the blastula-to-gastrula transition

Maximal-intensity projections were generated from the original z-stacks to produce two-dimensional representations of the epithelium. Images were contrast-adjusted to enhance membrane signal visualization. Cell segmentation was performed using Tissue Analyzer^117^, which was used to extract morphometric parameters including cell area and perimeter. Segmentation accuracy was subsequently verified by a second observer, and small false-positive objects, such as membrane-associated deposits or artifacts, were manually excluded from the analysis.

For each segmented cell, circularity was calculated as:

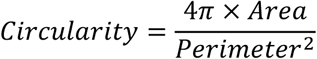

and cell shape index was calculated as:

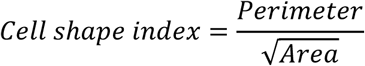

Average values for each field of view were then calculated and plotted using Python.

#### Morphometric analysis of tricellular zipping and vertex sliding

Cell segmentation and tracking was performed using Tissue Analyzer^117^. Segmentation, cell tracking, and junction tracking was verified and corrected manually. Cell area and junction lengths were extracted. Circularity was calculated as previously described, and junction length was normalized by the length of the junction at the completion of zipping. The angle analysis was performed by manually drawing the angles in FIJI for each junction and timepoint. For the circularity, junction length, and angles plots, the LOESS regression function for each group was plotted as a function of the percentage of time spent zipping using ggplot2 in R. Videos were cropped to only include the duration of zipping, so the percentage of time spent zipping was calculated by dividing the time by the duration of the video and converting to a percentage.

#### Quantification of tricellular zipping after cytokinesis

The end of cytokinesis was defined as the time point at which the cytokinetic furrow had fully ingressed and formed a higher-order vertex between the dividing cell and a neighboring cell. The onset of tricellular zipping was defined as the time point at which a tricellular vertex began moving away from the transient higher-order vertex formed after cytokinesis. The time interval between the end of cytokinesis and the onset of tricellular zipping was measured and plotted. The distances between the remodeling or stationary vertex and the midbody were measured in maximum-projection images using the line tool in Fiji. Line length was measured at each time point and normalized to its initial value, defined as measured length/initial length.

#### Quantification of de novo bicellular junction elongation

To measure the length of the *de novo* bicellular junction (BCJ), the straight-line tool in Fiji was used. On the maximum projected movies, the line was manually drawn from the anchor vertex to the remodeling vertex and the length of the line was measured in FiJi. For each time frame the line was manually adjusted between the anchor and remodeling vertex until completion of the zipping.

#### Quantification of E-cadherin and Myosin II intensity

E-cadherin-3xGFP and SF9-mCherry signals were measured in Fiji at the stationary vertex for each time frame, from the start to the end of the tricellular zipping process, using a circular ROI with a diameter of 2.90 µm, corresponding to 7 pixels. A Fiji macro was used to sequentially advance frames after each measurement. Similarly, E-cadherin-3xGFP and SF9-mCherry signals were measured at the remodeling vertex using a circular ROI with a diameter of 4.18 µm, corresponding to 14 pixels. A larger ROI was used for the remodeling vertex to capture the full signal around the actively remodeling vertex during zipping. Raw E-cadherin-3xGFP and SF9-mCherry intensities at the stationary and remodeling vertices were plotted for a single zipping event in Fig. 3C. For normalization, E-cadherin-3xGFP and SF9-mCherry signals were measured at a non-remodeling tricellular junction (TCJ) in the same field of view, but away from the zipping cells. A circular ROI with a diameter of either 2.90 µm, corresponding to 7 pixels, or 4.18 µm, corresponding to 14 pixels, was used to match the ROI size used for the stationary or remodeling vertex, respectively. E-cadherin and Myosin II intensities at the remodeling and stationary vertices were normalized as follows:

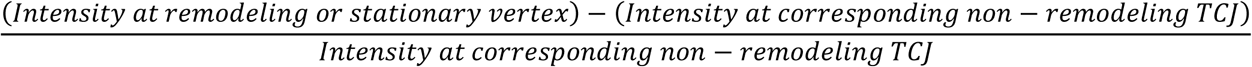

These normalized intensity values are plotted in Fig. 3F.

The maximum peak in the raw Myosin II and normalized Myosin II/E-cadherin intensity profile was determined in Python>3.8 using the find_peaks function from ‘SciPy’. Peaks that did not have any additional peaks within 2 min before or after were then manually selected for subsequent quantification. The local minima before and after the maximum peak were also determined using the same function on the inverted signal. The time point of the local minima before and after the maximum peak was identified and the slope of *de novo* BCJ was calculated for 2 min before and 2 min after (post) the recorded time points (Fig. 3E, G). For the control analysis without peaks (Fig. S2A, B), random points along the time axis were selected using two criteria: the points had to be at least ±2 min away from the start or end of the measurement and at least ±2 min away from the peaks identified in the previous step.

#### Quantification of ZO-1 and Myosin II intensity and change in junction length

Using a 2.90 µm (7 pixels) diameter circular ROI, BFP-ZO-1 signal was measured at the remodeling vertex and at the stationary vertex for each time frame from the start until end of the tricellular zipping process in FiJi. Using the same sized ROI, BFP-ZO-1 signal was also measured at a non-remodeling TCJ in the same field of view but away from the zipping cell as a reference. A FiJi macro was used to sequentially advance the frames after each measurement. The ZO-1 intensity at the remodeling and stationary vertex were normalized as follows: Intensity at remodeling or stationary vertex / Intensity at non-remodeling TCJ. Then, the difference was calculated: (Normalized intensity at remodeling vertex – normalized intensity at stationary vertex) for each time frame throughout the zipping process. This data is plotted in Fig. 3M. Similarly, using a 2.90 µm (7 pixels) and a 4.18 µm (14 pixels) diameter circular ROI, SF9-mCherry (Myosin II) signal was measured at the remodeling vertex and stationary vertex for each time frame. Then, the product of ZO-1 and Myosin II was calculated (Raw intensity of ZO-1 X Raw intensity of Myosin II) for both stationary and remodeling vertex at each time frame throughout the zipping process. This data is plotted in Fig. 3H.

The product of ZO-1 and Myosin II was plotted along with the length of the *de novo* BCJ for each zipping event. Maximum peak in ZO-1 X Myosin II was determined in python using scipy’s ‘find_peaks’ function with PEAK_SETTING = 0.3σ. The local minima before the maximum peak were also determined using the same function on the inverted signal. The time point of the local minima before the maximum peak was identified, and the slope of *de novo* BCJ was calculated for 2 min before^116^ and 2 min after (post) the recorded time point.

#### Quantification of FRAP recovery curves

For remodeling and stationary vertex FRAP experiments, the intensity of the bleached region of interest and a nearby reference tricellular vertex was quantified in all frames using a circular ROI (1.45 μm diameter). Each measurement was taken in triplicate and then averaged. The intensities were normalized by the following formula:

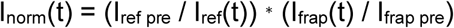

Where I_ref_ is the intensity of the reference junction, I_frap_ is the intensity of the bleached junction, (t) is the specific time point, and I_ref pre_ is the average intensity pre bleach. The normalized intensities were then constrained between 1 and 0 with the following formula:

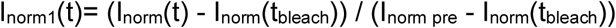

Where I_norm_ is the normalized value from the first formula, (t) is the specific time point, (t_bleach_) is the bleached time point, and I_norm pre_ is the average intensity pre-bleach time point. The I_norm1_ values were plotted over time using python and fit to a single exponential curve in order to extrapolate the mobile fraction and t_1/2_.

#### Quantification of calcium and F-actin intensity during tricellular zipping

To measure change in F-actin and calcium intensity during zipping, a rectangular ROI encompassing the long cell extension and the zipping junctions was drawn in FiJi. The ROI was overlayed on the maximum Z-projected images in Fiji, and mean intensity was measured. To accommodate for the movement in the live embryos, the ROI was manually moved to keep the zipping event in the center of the ROI for each time frame. The intensity of calcium was normalized as follows:

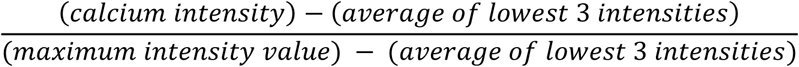

#### Quantification of pre– and post-calcium flash

To assess changes in *de novo* BCJ elongation before and after a calcium flash, individual calcium flashes were analyzed using a 4-minute window centered on the calcium flash. For each event, measurements of the *de novo* BCJ length were collected from 2 minutes before (–2 min) to 2 minutes after (+2 min) the flash, with the time of the flash defined as 0 min. Data were organized and plotted in Excel, and the *de novo* BCJ length was normalized by dividing each measurement by the measurement at +2 min. The normalized values were plotted over time to visualize changes in junction elongation relative to the calcium flash. To quantify the rate of junction elongation before and after the calcium flash, separate slopes were calculated for the pre-flash and post-flash periods in Excel. The measurement at 0 min, corresponding to the calcium flash, was excluded from the slope calculation. Slopes were calculated using the normalized *de novo* BCJ measurements from 2 minutes before to the flash and from the flash to 2 minutes after. The resulting pre– and post-flash slope values were used to plot a spaghetti plot in R.

#### Calcium and F-actin cross-correlation analysis

F-actin and calcium cross-correlation analysis was performed on the rates of change in calcium and F-actin intensities within the cell extension using a previously published cross-correlation code in MATLAB^118^.

#### Quantification of calcium, F-actin, and ZO-1 intensity in calcium uncaging experiment

To measure change in F-actin and calcium intensity during zipping, a rectangular ROI encompassing the long extension and the zipping junctions was drawn in FiJi. The ROI was overlayed on the maximum Z-projected images in Fiji, and mean intensity was measured for each time frame 2 min before and 2 min after the stimulation. To accommodate for the movement in the live embryos, the ROI was manually moved to keep the zipping event in the center of the ROI for each time frame. For normalization, a similar ROI was used to measure calcium intensity in the cytoplasm and F-actin intensity at a BCJ of another cell in the same field of view. The intensity of calcium was normalized in two steps:

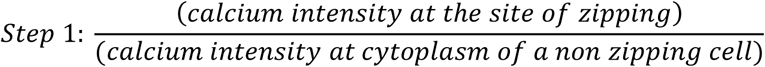

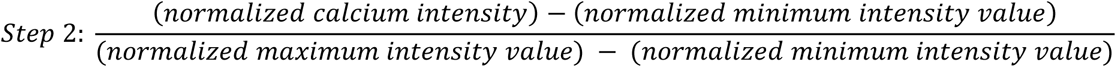

The intensity of F-actin was normalized as follows:

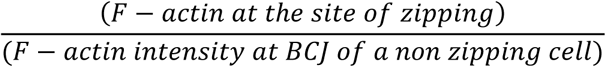

Using a 10 pixel diameter circular ROI, mRFP-ZO-1 signal was measured at the remodeling vertex, 2 min before and 2 min after the stimulation in FiJi. Using the same sized ROI, mRFP-ZO-1 signal was also measured at a BCJ in the same field of view but away from zipping cell as a reference. The intensity of ZO-1 was normalized as follows:

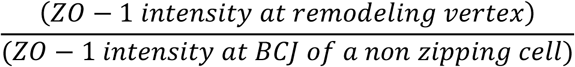

#### Quantification of calcium flash events in Drosophila

Time series videos were processed using Fiji. Apical views of embryos were obtained through maximum projections over a full Z-stack. Calcium flashes were manually counted and then validated through measuring the fold change between the membrane-bound GCaMP8s intensity before, during, and after calcium flash events and a ubiquitously expressed fluorescent membrane marker, gap43::mCherry7, to confirm that increases in GCaMP8s intensity are a result of calcium influx. To measure cell membrane intensity profiles surrounding a flash, linear ROIs were drawn across the cell and signal intensity was recorded one frame before, during, and after the flash for both the GCaMP8s and gap43::mCherry. Maximum intensity values at the membrane for each cell were averaged together for a singular average intensity value for each timepoint per channel. The calcium signal intensity was then normalized to the gap43::mCherry signal intensity, which allowed for fold change comparison across several replicates. Quantitative analysis and Wilcoxon signed rank tests were performed using GraphPad Prism 11.0.0 (GraphPad Software, Inc).

#### Identification of calcium events and bulk quantification of physical cell properties

Full-time series videos of *Drosophila* gastrulation were maximum Z-projected in Fiji, and cell segmentation masks were generated in Cellpose-SAM using the SAM-ViTL backbone as a model^80^. Due to the dynamic nature of tissue morphogenesis, cell masks were manually corrected across frames and interpolated to achieve complete segmentation of the embryo over the course of the timeseries. An example is displayed in Fig. S5B.

Bulk quantification of cell properties using those masks was done with CellProfiler using a custom pipeline^119^. The TrackObjects function was used to ensure continuity of cellIDs for each frame, and X and Y coordinates for each cell were also recorded. Cell properties including area, major and minor axis length, circularity, and GCaMP intensity were all measured and recorded for each individual cell across the timeseries.

A custom Python script was written to identify calcium flashes in cells using the data gathered from the CellProfiler pipeline. Calcium flash events were defined as having a ΔF/F local baseline threshold value of at least 0.3 and an event peak ΔF/F value of at least 0.35 for a maximum of 60 seconds. ΔF/F is calculated by the following formula: 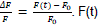 F(t) is defined as the GCaMP8s intensity value for the cell at time (t), and F_0_ is the baseline intensity which is measured using a rolling local percentile of the intensity for a cell over time and has been set to be the 20th percentile within the minimum rolling window of about 22 seconds. This baseline accommodates the dynamic GCaMP8s signal and minimizes false positives along with recording calcium events in dimmer regions of the embryo. 76/96 of events detected from the parameters set in the Python script were previously identified manually through visual inspection in Fiji. Data from identified calcium flash events were then peak-aligned at t=0 and observed within a 4-minute timeframe (± 2 minutes). All quantitative analysis and Wilcoxon signed rank tests were performed using GraphPad Prism 11.0.0.

### Video editing

Videos were initially generated and concatenated using Fiji 2.16.0/1.5p. Overlay animations were added using Adobe Premiere Pro 25.5.

### Data availability

The datasets generated during and/or analyzed during the current study are available from the corresponding authors on reasonable request.

## Acknowledgments

We thank all the members of the Miller and Martin labs for helpful discussions and feedback on this research. We thank the He lab and Littleton lab for the *Drosophila* reagents and the Bement lab for 3xGFP-ROCK2 K106A construct. We would like to express our gratitude to Janis F. Miroll for assistance with statistical analysis and plotting with Python. We acknowledge the University of Michigan BRCF Microscopy Core (RRID: SCR 026722) for training and use of the Zeiss LSM 980 with Airyscan2 confocal microscope. We are grateful to the NIH-funded National *Xenopus* Resource (RRID: SCR_013731) and *Xenbase* (RRID: SCR_003280) for public support of *Xenopus* research.

## Author contributions

B.A., A.L.M., A.C.M., and S.B. conceived the study. B.A., A.S., and A.G.L-F. collected and analyzed data in *Xenopus laevis*. X.O.Z and M.P. collected data and analyzed data in *Drosophila*. B.A and T.R.H. collected mouse data. K.K.T. analyzed zipping morphometric data. S.B. developed the theory and performed simulation. B.A., X.O.Z., and S.B. wrote the original draft. All authors revised the paper and approved the final version. B.A., A.L.M. and A.C.M. supervised the study.

## Funding

This work was funded by NIH R35 GM153204 to A.L.M, NIH R35 GM144115 to A.C.M., NIH R35 GM143042 to S.B., and lab startup funds from the University of Michigan Department of Molecular, Cellular, and Developmental Biology to T.R.H. B.A. acknowledges the support of University of Michigan Mary Sue Coleman Fellowship, Barbour Scholarship, and Rackham candidate research grant.

## Declaration of interests

The authors declare no competing financial interests.

## Declaration of generative AI and AI-assisted technologies

During analysis of the data, the authors used Claude model Opus 4 & 5 to draft and debug code to analyze, normalize, and plot the quantified data. After using the tool, the authors reviewed and edited the code as needed and take full responsibility for the publication’s content.

## Notes

### Competing Interest Statement

The authors have declared no competing interest.

