## Supplementary Information for "Calcium and actomyosin pulses drive tricellular zipping to remodel epithelial cell-cell junctions and facilitate cell packing"

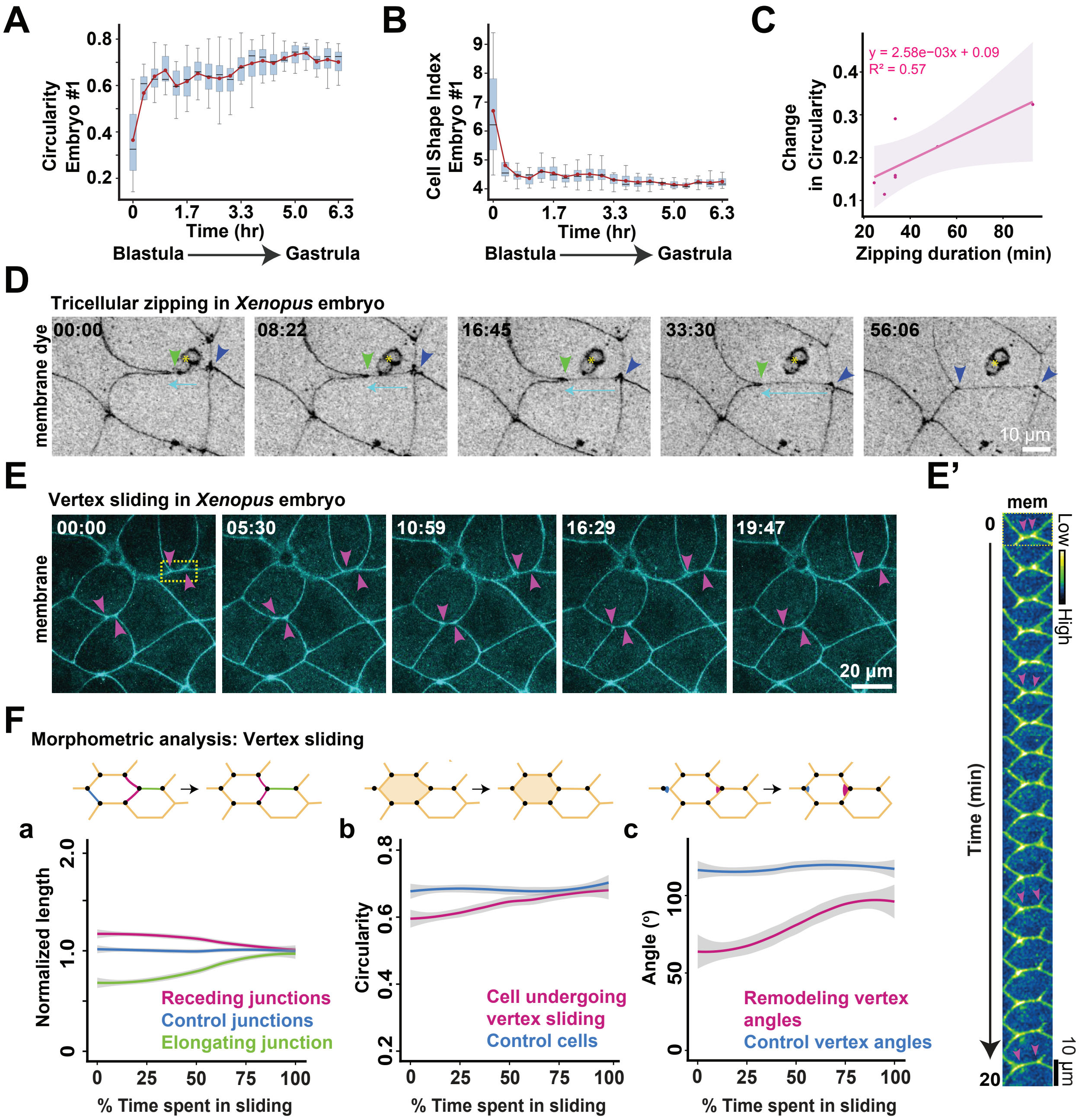
A
Circularity Embryo #1
Time (hr)
Blastula
Gastrula
B
Cell Shape Index Embryo #1
C
y = 2.58e−03x + 0.09
R² = 0.57
Change in Circularity
Zipping duration (min)
D
Tricellular zipping in *Xenopus* embryo
membrane dye
00:00
08:22
16:45
33:30
56:06
10 µm
E
Vertex sliding in *Xenopus* embryo
membrane
00:00
05:30
10:59
16:29
19:47
20 µm
E'
mem
Low
High
Time (min)
10 µm
F
Morphometric analysis: Vertex sliding
a
Normalized length
Receding junctions
Control junctions
Elongating junction
% Time spent in sliding
b
Circularity
Cell undergoing vertex sliding
Control cells
c
Angle (°)
Remodeling vertex angles
Control vertex angles

**Supplementary Figure 1. Tricellular zipping facilitates regularization of epithelial cell packing in embryos and differs from vertex sliding.**

**(A-B)** Quantification of cell circularity (A) and cell shape index (B) for cells within the field of view during the blastula-to-gastrula transition in a single embryo. The solid red line represents the mean. In the box plots, the black line indicates the median, and whiskers indicate the minimum and maximum values at each time point. **(C)** Correlation plot showing the relationship between the change in circularity, measured before zipping and after zipping completes, of receding cells and the time spent in zipping. $n$ = 7 zipping events, 3 independent experiments. **(D)** Time-lapse confocal images of tricellular zipping in a *Xenopus laevis* embryo with cell membranes labeled using CellMask dye. Green arrowheads indicate the remodeling vertex, blue arrowheads indicate the stationary vertex, cyan arrows indicate the direction of zipping, and yellow asterisk indicates dye labeling a midbody trapped between the embryo and vitelline envelope from an earlier cytokinesis. Time is shown as min:sec from the onset of zipping. Scale bar, 10 μm. **(E)** Time-lapse confocal images of vertex sliding in a *Xenopus laevis* embryo expressing a membrane probe (mem-BFP). Magenta arrowheads indicate sliding vertices. Time is shown as min:sec. Scale bar, 20 μm. **(E′)** Kymograph generated from the yellow boxed region shown in (E). Green Blue Fire LUT. **(F)** Quantification of morphometric changes during vertex sliding. Schematics illustrate the vertex sliding process and the measured parameters. (a) Vertex-to-vertex lengths of junctions were measured over time and normalized to their respective initial lengths at the onset of vertex sliding. (b) Circularity was measured for cells involved in vertex sliding (magenta) and for control cells within the same field of view (blue). (c) The angles of the remodeling vertex (magenta) and control vertices (blue) were measured during vertex sliding. Nonlinear regression across multiple vertex sliding events is plotted over a normalized time axis representing the percentage of total time spent sliding. Solid lines indicate the fitted mean, and shaded regions represent the 95% confidence interval. For (a), $n$ = 10 elongating junctions, 19 receding junctions, and 27 control junctions, 9 embryos, 3 experiments. For (b), $n$ = 10 sliding cells and 20 control cells, 9 embryos, 3 experiments. For (c), $n$ = 11 remodeling vertices and 21 control vertices, 9 embryos, 3 experiments.

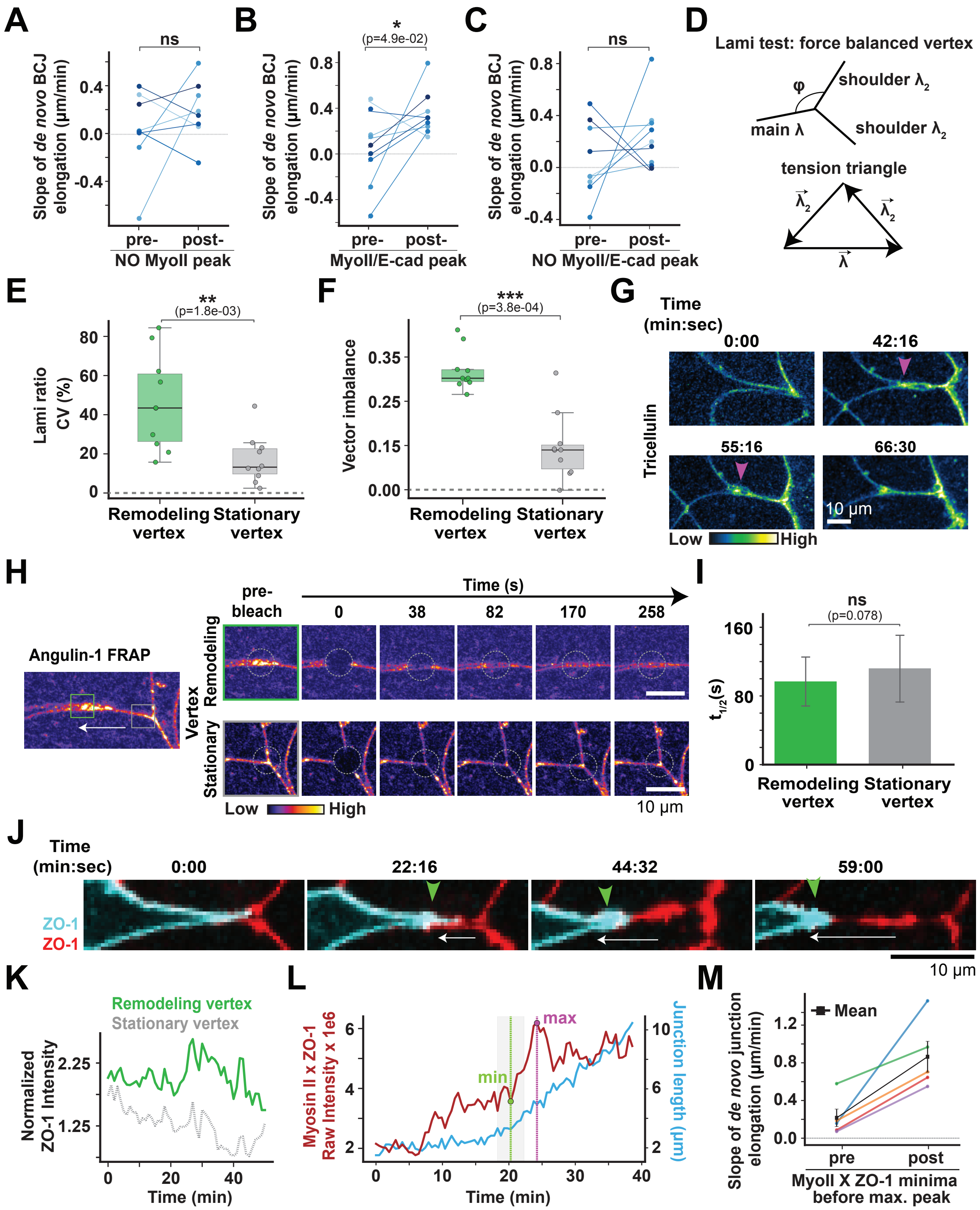

A
ns
Slope of de novo BCJ elongation (µm/min)
NO MyoII peak
pre-
post-
B
*
(p=4.9e-02)
MyoII/E-cad peak
C
NO MyoII/E-cad peak
D
Lami test: force balanced vertex
φ
shoulder λ2
main λ
shoulder λ2
tension triangle
E
**
(p=1.8e-03)
Lami ratio CV (%)
Remodeling vertex
Stationary vertex
F
***
(p=3.8e-04)
Vector imbalance
G
Time (min:sec)
0:00
42:16
55:16
66:30
Tricellulin
10 µm
Low
High
H
Angulin-1 FRAP
pre-bleach
Time (s)
0
38
82
170
258
Vertex
Remodeling
Stationary
Low
High
10 µm
I
ns
(p=0.078)
t1/2(s)
Remodeling vertex
Stationary vertex
J
Time (min:sec)
0:00
22:16
44:32
59:00
ZO-1
ZO-1
10 µm
K
Remodeling vertex
Stationary vertex
Normalized ZO-1 Intensity
Time (min)
L
max
min
Myosin II x ZO-1 Raw Intensity x 1e6
Junction length (µm)
Time (min)
M
Mean
Slope of de novo junction elongation (µm/min)
pre
post
MyoII X ZO-1 minima before max. peak

**Supplementary Figure 2. Tension asymmetry and differential force transmission drives displacement of the remodeling vertex**

**(A)** Quantification of paired slopes of *de novo* BCJ elongation for a region where there was NO peak in Myosin II intensity from individual zipping events (control for Fig. 3E and F). $n$ = 9 zipping events, 7 embryos, 4 experiments. **(B)** Quantification of paired slopes of *de novo* BCJ elongation, before and after a peak in the Myosin II/E-cadherin ratio from individual zipping events (see Fig. 3G). **(C)** Quantification of paired slopes of *de novo* BCJ elongation for a region where there was NO peak in Myosin II/E-cadherin ratio from individual zipping events (control for Fig. S2B). For (B) and (C), $n$ = 9 zipping events, 9 embryos, 4 experiments. Statistical significance for (A), (B) and (C) were determined using a paired t-test; $*p \leq 0.5$ and ns, $p > 0.5$. **(D)** Lami test to determine whether the stationary vertex is force balanced. Top: three junctions meet with interior angles $\varphi$; equilibrium requires the tension vectors to close into a triangle (bottom). The Lami relation (see methods) tests, from measured angles and myosin, whether the arcs are load-bearing and balanced. **(E)** Box plot showing Lami ratio coefficient of variation (CV) % of remodeling and stationary vertices. **(F)** Box plot showing vector imbalance of remodeling and stationary vertices during zipping. For both (E) and (F), box plot shows the median (solid line), the box shows the $25^{th}$ to $75^{th}$ percentile, and the whiskers represent the min and max values. $n$ = 10 stationary and remodeling vertices from 10 independent zipping events; 7 embryos, 6 experiments. Statistical significance was determined using a paired-$t$ test; $**p \leq 0.01$. Gray dashed line at 0 represents a perfectly stable vertex. **(G)** Time-lapse confocal images of tricellular zipping in *Xenopus* embryos expressing GFP-Tricellulin-1 (Green Fire Blue LUT). Tricellulin accumulates at the remodeling vertex (magenta arrowheads) and moves in the direction of zipping. Time is shown as min:sec. Scale bar, 10 µm. **(H)** Montage of Angulin-1-3xGFP (FIRE LUT) FRAP of a remodeling and a stationary vertex during zipping. Dashed circle indicates photobleached region. Time is shown as sec. Scale bar, 10 µm. **(I)** $t_{1/2}$ calculated from (Fig. 3I). Statistics, Mann-Whitney U test; $p > 0.05$ (ns); error bars indicate SEM. **(J)** Time-lapse confocal images of tricellular zipping in *Xenopus* embryos mosaically expressing two colors of tagged ZO-1: TagBFP2-ZO-1 (cyan) and ZO-1-RFP (red). ZO-1 accumulates at the remodeling vertex in the receding cell (green arrowheads) and moves in the direction of zipping (white arrow). Time is shown as min:sec. Scale bar, 10 µm. **(K)** Representative intensity profile of ZO-1 at the remodeling (green) and stationary (gray) vertices during a single zipping event. **(L)** Representative plot showing the product of raw intensity of Myosin II X ZO-1 at the remodeling vertex (red) along with length of the *de novo* BCJ (blue) during a single zipping event. Maximum peak in Myosin II X ZO-1 is identified (magenta dotted line), and the local minima before the peak is identified (green dotted line), and these values are used for slope analysis (see Fig. S2M). **(M)** Quantification of paired slope of *de novo* BCJ length pre- and post- the local minima before the maximum peak in Myosin II X ZO-1 product at the remodeling vertex during zipping. Black line shows mean. *n* = 5 zipping events, 5 embryos, 5 experiments.

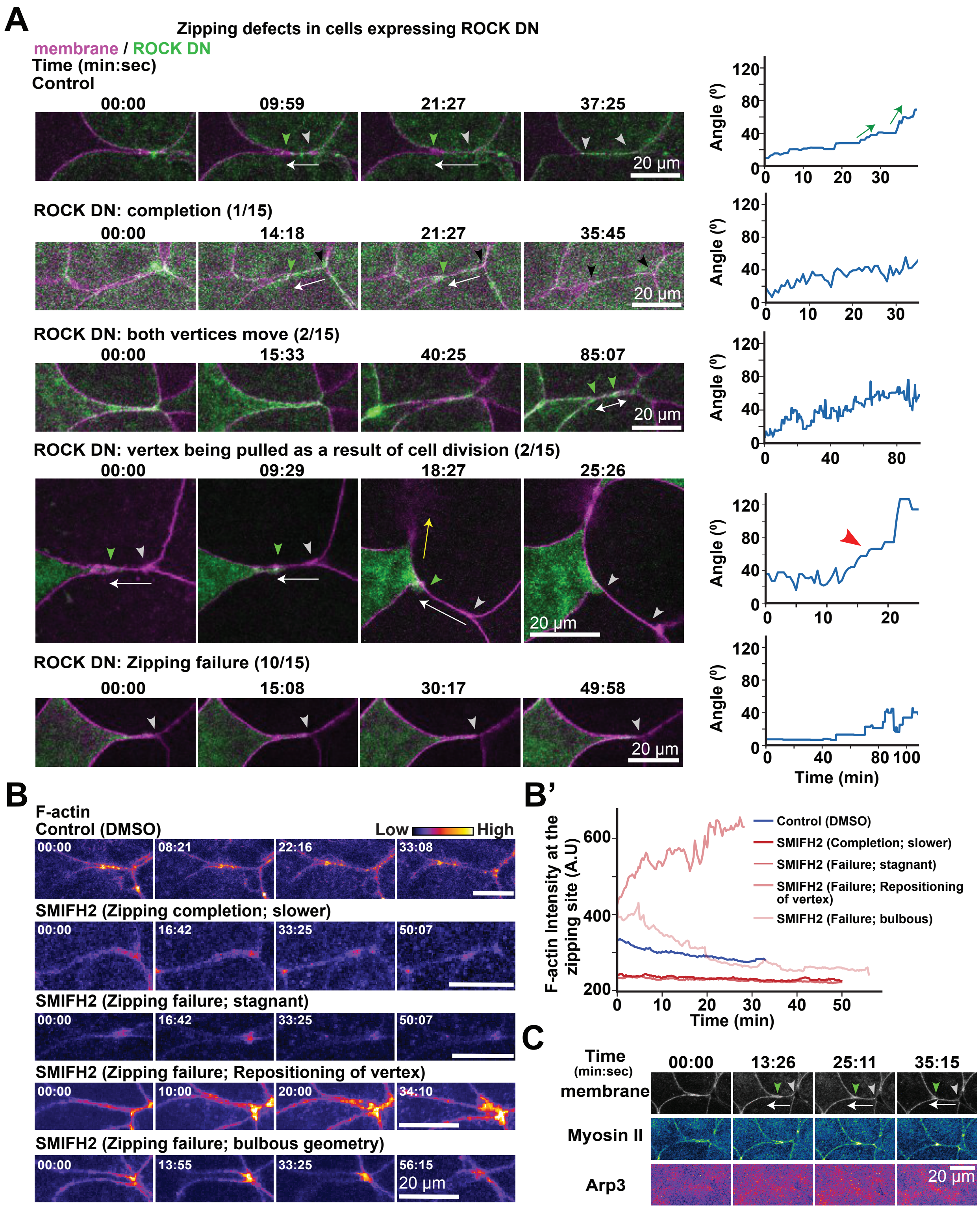

A
Zipping defects in cells expressing ROCK DN
membrane / ROCK DN
Time (min:sec)
Control
00:00
09:59
21:27
37:25
20 µm
ROCK DN: completion (1/15)
00:00
14:18
21:27
35:45
ROCK DN: both vertices move (2/15)
00:00
15:33
40:25
85:07
ROCK DN: vertex being pulled as a result of cell division (2/15)
00:00
09:29
18:27
25:26
ROCK DN: Zipping failure (10/15)
00:00
15:08
30:17
49:58
Angle (º)
Time (min)
B
F-actin
Control (DMSO)
Low
High
00:00
08:21
22:16
33:08
SMIFH2 (Zipping completion; slower)
16:42
33:25
50:07
SMIFH2 (Zipping failure; stagnant)
SMIFH2 (Zipping failure; Repositioning of vertex)
10:00
20:00
34:10
SMIFH2 (Zipping failure; bulbous geometry)
13:55
56:15
B'
F-actin Intensity at the zipping site (A.U)
Time (min)
Control (DMSO)
SMIFH2 (Completion; slower)
SMIFH2 (Failure; stagnant)
SMIFH2 (Failure; Repositioning of vertex)
SMIFH2 (Failure; bulbous)
C
Time (min:sec)
00:00
13:26
25:11
35:15
membrane
Myosin II
Arp3

**Supplementary Figure 3. Localized actomyosin dynamics drive tricellular zipping.**

**(A)** Representative examples of different kinds of zipping defects observed in cells expressing ROCK DN. Green arrowheads show the remodeling vertex, gray / black arrowheads show the stationary vertex, yellow arrow shows pull by the cytokinetic furrow of a dividing cell, double headed white arrow shows movement of both vertices, and single headed white arrows show the direction of zipping. Time is shown as min:sec. Scale bars, 20 μm. (Right) the corresponding quantification of angle of the remodeling vertex during the tricellular zipping. Green arrows show stepwise increase in angle during zipping, and red arrowhead shows sudden increase in angle due to neighboring cell division forces. **(B)** Representative examples of zipping defects observed in embryos expressing probe for F-actin (LifeAct-miRFP703, FIRE LUT) that were treated with either 0.2% DMSO (vehicle control) or 100 μM SMIFH2. Time is shown as min:sec. Scale bars, 20 μm. **(B')** Quantification of F-actin intensity at the zipping site for the examples shown in Fig. S3B. **(C)** Representative time-lapse confocal images of tricellular zipping in a *Xenopus laevis* embryo expressing a membrane probe (mem-BFP, grayscale), Myosin II probe (SF9-mCherry, Green Fire Blue LUT), and Arp3-mNeon (FIRE LUT). Green arrowheads indicate the remodeling vertex, gray arrowheads indicate the stationary vertex, and white arrows indicate the direction of zipping. Time is shown as min:sec from the onset of zipping. Scale bar, 20 μm.

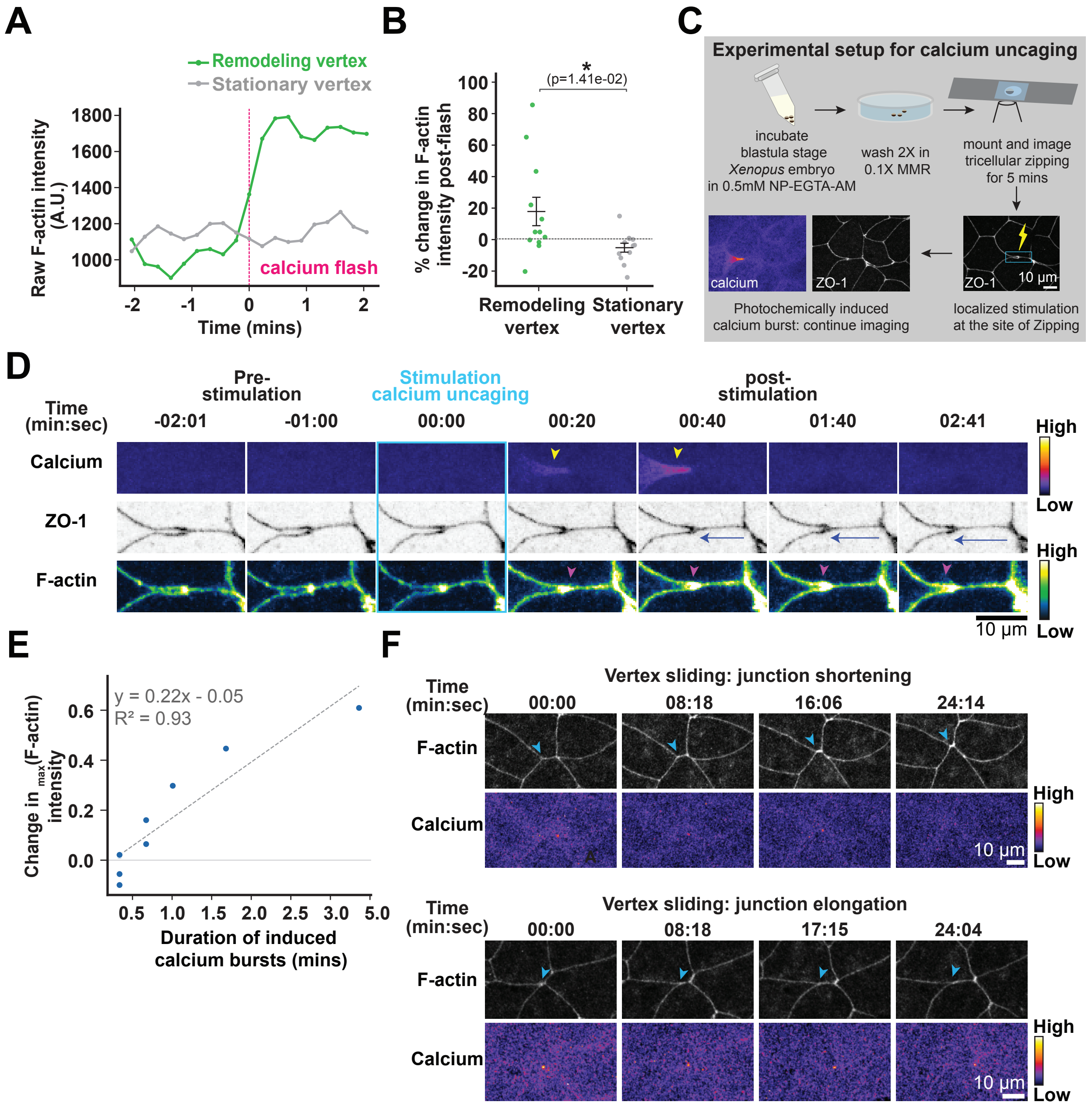

A
Remodeling vertex
Stationary vertex
Raw F-actin intensity (A.U.)
calcium flash
Time (mins)
B
(p=1.41e-02)
% change in F-actin intensity post-flash
Remodeling vertex
Stationary vertex
C
Experimental setup for calcium uncaging
incubate blastula stage Xenopus embryo in 0.5mM NP-EGTA-AM
wash 2X in 0.1X MMR
mount and image tricellular zipping for 5 mins
calcium
ZO-1
10 µm
Photochemically induced calcium burst: continue imaging
localized stimulation at the site of Zipping
D
Pre-stimulation
Stimulation calcium uncaging
post-stimulation
Time (min:sec)
-02:01
-01:00
00:00
00:20
00:40
01:40
02:41
Calcium
ZO-1
F-actin
High
Low
10 µm
E
y = 0.22x - 0.05
R² = 0.93
Change in max(F-actin) intensity
Duration of induced calcium bursts (mins)
F
Vertex sliding: junction shortening
Time (min:sec)
00:00
08:18
16:06
24:14
F-actin
Calcium
High
Low
10 µm
Vertex sliding: junction elongation
Time (min:sec)
00:00
08:18
17:15
24:04
F-actin
Calcium
High
Low
10 µm

**Supplementary Figure 4. Calcium flashes are associated with tricellular zipping.**

**(A)** Representative intensity profile of F-actin at the remodeling (green) and stationary (gray) vertices pre- (2 min before) and post- (2 min after) a calcium flash during tricellular zipping (dotted magenta line marks the time point corresponding to maximum intensity of calcium). **(B)** Quantification showing % change in mean F-actin intensity at the remodeling and stationary vertex 2 min before and after the calcium flash during zipping. n=12 remodeling and stationary vertices corresponding to 12 zipping events, 11 embryos, 8 independent experiments. Statistical significance was determined using Mann-Whitney U test; $^*p \leq 0.05$. **(C)** Experimental setup for localized calcium uncaging experiment. **(D)** Time-lapse confocal images of tricellular zipping in a *Xenopus laevis* embryo before and after induced calcium bursts, labeled with probes for calcium (GCaMP6s), ZO-1 (mRFP-ZO-1), and F-actin (LifeAct-miRFP703). Localized stimulation to induce calcium burst was performed at time point 0 shown by the blue box. Yellow arrowheads indicate the increase in calcium, blue arrows indicate increased rate of zipping, and magenta arrowheads indicate increase in F-actin right after the stimulation. Time is shown as min:sec. Scale bar, 10 μm. **(E)** Correlation plot showing the relationship between the change in maximum F-actin intensity at the site of zipping, measured during 2 min prior and 2 min after stimulation, and the duration of induced calcium bursts. $n$ = 10 zipping events, 10 embryos, 5 independent experiments. **(F)** Time-lapse montage of vertex sliding (blue arrowheads) in a *Xenopus laevis* embryo labeled for F-actin (LifeAct-miRFP703, grayscale) and calcium (GCaMP6m, FIRE LUT). Time is shown as min:sec. Scale bars, 10 μm.

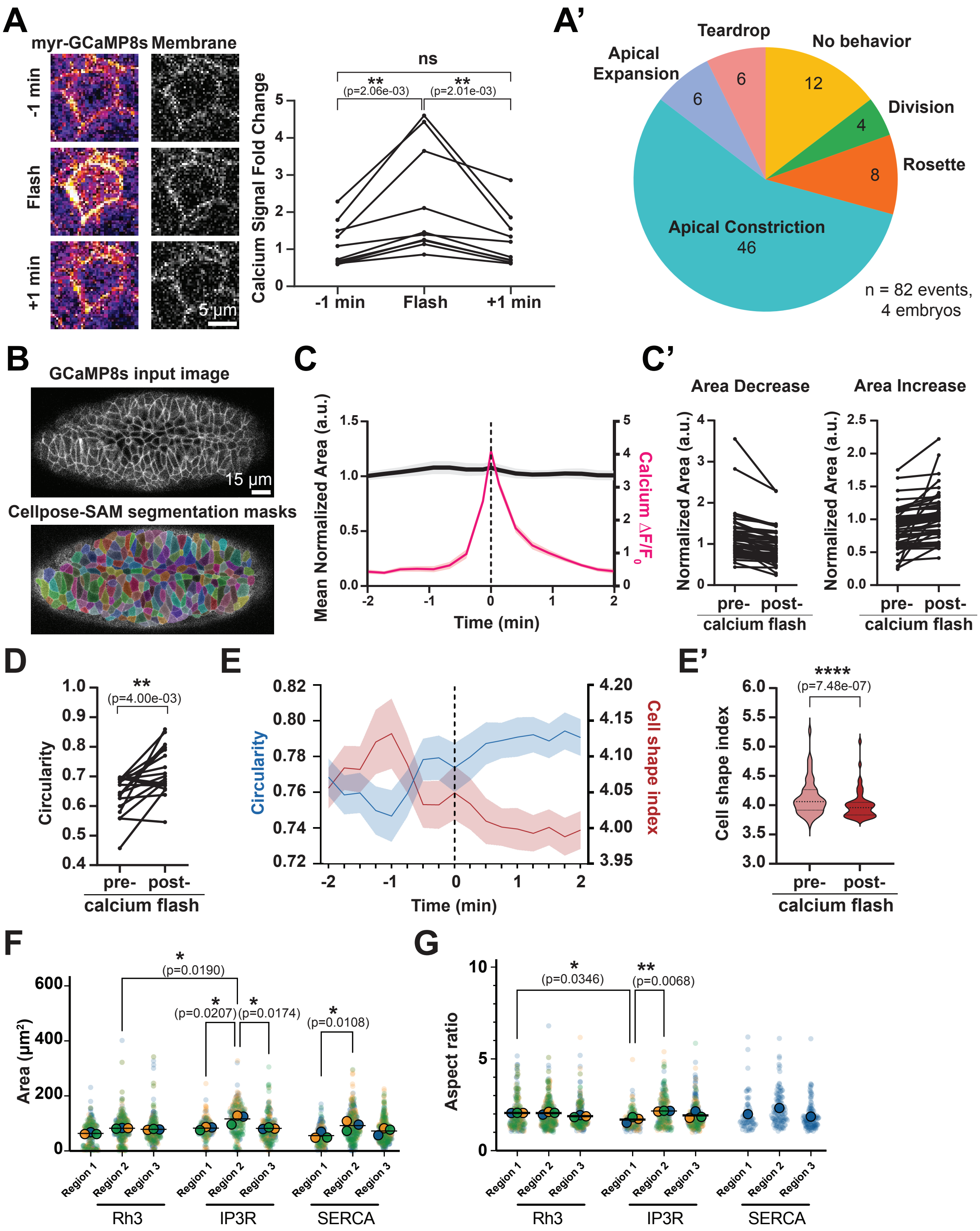

A
myr-GCaMP8s
Membrane
-1 min
Flash
+1 min
5 µm
Calcium Signal Fold Change
ns
**
(p=2.06e-03)
**
(p=2.01e-03)
-1 min
Flash
+1 min
A'
Teardrop
No behavior
Apical Expansion
Division
Rosette
Apical Constriction
46
n = 82 events, 4 embryos
B
GCaMP8s input image
15 µm
Cellpose-SAM segmentation masks
C
Mean Normalized Area (a.u.)
Calcium ΔF/F0
Time (min)
C'
Area Decrease
Area Increase
Normalized Area (a.u.)
pre- post-
calcium flash
D
**
(p=4.00e-03)
Circularity
pre- post-
calcium flash
E
Circularity
Cell shape index
Time (min)
E'
****
(p=7.48e-07)
Cell shape index
pre- post-
calcium flash
F
Area (µm²)
*
(p=0.0190)
*
(p=0.0207)
*
(p=0.0174)
*
(p=0.0108)
Region 1
Region 2
Region 3
Rh3
IP3R
SERCA
G
Aspect ratio
*
(p=0.0346)
**
(p=0.0068)
Region 1
Region 2
Region 3
Rh3
IP3R
SERCA

**Supplementary Figure 5. Calcium flashes are correlated with cell shape changes during *Drosophila* gastrulation.**

**(A)** Timelapse confocal images of a representative calcium flash in a *Drosophila melanogaster* embryo expressing a membrane-bound probe for cytoplasmic calcium (myr-GCamP8s, FIRE LUT) and and a membrane marker (gap43-mCherry7, grayscale). Scale bar, 5 μm. Quantification of calcium intensity from flash events (right). Calcium signal was normalized against the gap43 membrane signal. $n = 10$ events, 4 embryos. Statistical significance was determined using a Wilcoxon matched-pairs signed rank test, $^{**}p < 0.01$**.** **(A')** Qualitative assessment of cell shape change behaviors associated with calcium flashes. Behaviors were assigned to each flash event based on shape and area changes ± 2 minutes from calcium flash; one primary behavior was assigned per calcium flash. $n = 82$ events, 4 embryos. **(B)** Example image and resulting segmentation masks from Cellpose-SAM model for bulk-segmentation of *Drosophila* calcium time-lapse movies. Scale bar, 15 μm. **(C)** Quantification of the mean normalized area (gray) and $\Delta F/F0$ (red) during calcium flash events. All event peaks are aligned at $t = 0$ min (dotted line). Mean (solid line) ± SEM (shading). $n = 80$ events, 5 embryos. **(C')** Quantification of changes in apical area 1 min before and 1 min after a calcium flash, sorted into 'Area Decrease' (left) and 'Area Increase' (right) bins. For 'Area Decrease,' $n = 47$ events, 5 embryos. For 'Area Increase,' $n = 49$ events, 5 embryos. **(D)** Quantification of changes in cell circularity of the least-circular cells (initial circularity < 0.7) during calcium flash events. $n = 18$ events, 4 embryos. Statistical significance was determined using a Wilcoxon matched-pairs signed rank test, $^{**}p < 0.01$. **(E)** Quantification of mean cell circularity (blue) and cell shape index (red) during calcium flash events for cells that increase in circularity. All event peaks are aligned at $t = 0$ min (dotted line). Mean (solid line) ± SEM (shading). $n = 41$ events, 5 embryos**.** **(E')** Quantification of changes in cell shape index for cells that increase in circularity during calcium flash events. $n = 50$ events, 5 embryos. Statistical significance was determined using a Wilcoxon matched-pairs signed rank test, $^{****}p < 0.0001$. **(F)** Quantification of cell area by embryonic region in control, IP3R, and SERCA RNAi knockdown embryos at stage 3. $N = 3$ embryos per genotype, with a minimum of 46 cells quantified per embryo. Statistical significance was determined using Tukey's multiple comparisons test; $^{*}p \leq 0.05$. **(G)** Quantification of aspect ratio by embryonic region in control, IP3R, and SERCA RNAi knockdown embryos at stage 3. $N = 3$ embryos per genotype, with a minimum of 46 cells quantified per embryo. Statistical significance was determined using Tukey's multiple comparisons test; $^{*}p \leq 0.05$, $^{**}p \leq 0.01$.

**Mechanical model of tricellular zipping**

Here we describe the mechanical framework used to interpret the asymmetric motion of the two vertices bounding the bicellular junction during tricellular zipping (Fig. 3A), the two hypotheses tested against the molecular data (Fig. 3B–L, S2), the geometric test of force balance at the stationary vertex (Fig. S2D–F), and the calcium-gated dynamical model used to simulate the response of the junction to an induced calcium flash (Fig. 6H–K). The model presented is minimal and phenomenological; the goal is to organize the observations and to make the logical structure of the inferences explicit. All comparisons with experiment are qualitative.

Force-balance framework – We index the remodeling vertex by $i = 1$ and the stationary vertex by $i = 2$. Each vertex is pulled by the tensions of the junctions meeting there. The two shoulder junctions of vertex $i$ carry tension $\lambda_i$ and make half-angle $\theta_i$ with the zipping axis, and the bicellular junction carries tension $\lambda$ and opposes advance (Fig. 3A). The net junctional force on vertex $i$ along the zipping axis is

$$F_i = 2\lambda_i cos\theta_i - \lambda. \qquad (S1)$$

Vertex motion in the embryonic epithelium is overdamped, so velocity is proportional to the net force that is actually delivered to the vertex. We write this as

$$\gamma_i v_i = \alpha_i F_i, \qquad (S2)$$

where $\gamma_i$ is a friction coefficient and $\alpha_i \in [0,1]$ is the force transmission efficiency, which represents the fraction of the junctional force generated by the actomyosin network that is mechanically coupled to the vertex. We interpret the data using Eq. (S2), which is general in scope. A vertex moves when the transmitted junctional forces do not cancel. Conversely, a vertex is stationary when the forces balance, or when nothing is transmitted.

*Tension asymmetry* – Junctional tension is set by actomyosin, and we take $\lambda_i$ to increase with local Myosin II. Myosin II is enriched at the remodeling vertex (Fig. 3B–D) and organizes into a horseshoe-shaped pattern that moves with the remodeling vertex (Fig. 4A), so

$$\lambda_1 > \lambda_2 \quad \Rightarrow \quad F_1 > F_2. \qquad (S3)$$

This asymmetry provides the net force that displaces the remodeling vertex. Local peaks in Myosin II at the remodeling vertex precede the transient increases in the rate of *de novo* BCJ elongation (Fig. 3E–F), consistent with $v_1$ tracking $F_1$ through Eq. (S2).

**Two hypotheses for the immobility of the stationary vertex** – The force-balance framework doesn't answer why the stationary vertex feels no net force although it is embedded in the same tension network. Eq. (S2) suggests two hypotheses, which make opposite predictions about protein localization.

Friction hypothesis – The stationary vertex is subject to active force but is held in place by a large resistance to motion, $\gamma_2 \gg \gamma_1$, supplied by adhesive anchoring. This predicts enrichment of the adhesion machinery, such as E-cadherin, at the stationary vertex.

Clutch hypothesis – The stationary vertex is weakly coupled to the contractile force generating machinery, $\alpha_2 \ll \alpha_1$, so that active force is not efficiently transmitted to it. The analogy is the molecular clutch that couples actomyosin to adhesions in migrating cells[1, 2]. This predicts enrichment of linkers that couple actomyosin to junctions, such as ZO-1, at the remodeling vertex, where force is being transmitted.

**The friction hypothesis is not supported by data** – Every marker of adhesion machinery that we examined contradicts the friction hypothesis. First, E-cadherin is enriched at the stationary vertex in only 7/14 zipping events (Fig. 3B–D). The amount of adhesion does not determine which vertex is stationary. Second, ZO-1, a candidate anchoring element, is consistently higher at the remodeling vertex (Fig. 3K–L, S2J–K). A drag element that decorates the faster vertex is the opposite of what $\gamma_2 \gg \gamma_1$ requires. By FRAP, Angulin-1 has the higher mobile fraction at the stationary vertex, $0.7 \pm 0.06$ versus $0.4 \pm 0.05$ (Fig. 3I–J). The stationary vertex is thus a more fluid structure that is turning over fast, whereas a vertex held by a high-drag anchor would be expected to be the less dynamic one. The ratio of Myosin II to E-cadherin intensity, which we had originally treated as a proxy for tension-to-friction ratio, shows peaks at the remodeling vertex that precede the accelerated elongation (Fig. 3G, S2B–C). Because the same result is obtained from Myosin II alone (Fig. 3F) and E-cadherin is not asymmetric, we interpret this ratio as a measure of contractility rather than as evidence for a drag term.

**The stationary vertex is close to mechanical equilibrium** – If the stationary vertex is not anchored by drag, the alternative is that the forces acting on it cancel, so that it sits at mechanical equilibrium. This is testable from cell geometry alone. At a vertex where the three junctions meet, equilibrium requires the three tension vectors to sum to zero. This is equivalent to Lami's theorem: if $\alpha_a, \alpha_b, \alpha_c$ are the angles opposite junctions $a, b, c$, (with tensions $T_a$, $T_b$, and $T_c$, respectively) then (Fig. S2D)

$$\frac{T_a}{sin\alpha_a} = \frac{T_b}{sin\alpha_b} = \frac{T_c}{sin\alpha_c}. \qquad (S4)$$

We use mean Myosin II intensity along each junction, $m_j$, as a relative tension proxy, $T_j \propto m_j$. Eq. (S4) involves only tension ratios, so the unknown proportionality constant drops out. We computed two per-vertex measures of departure from balance (Fig. S2D–F). First, we calculated the coefficient of variation of $r_j = \frac{m_j}{sin\alpha_j}$ across the junctions at a vertex,

$$CV = \frac{std(r_j)}{mean(r_j)}. \qquad (S5)$$

At equilibrium, the ratios are equal and $CV = 0$. Second, we computed the vector imbalance at the junctions. Each junction is assigned a tension vector of magnitude $m_j$ directed along the junction, with directions $\hat{n}_j$ obtained by walking the measured sector angles around the vertex. The imbalance is the magnitude of the net uncancelled pull as a fraction of the total tension,

$$I = \frac{\| \sum_j \quad m_j\hat{n}_j \|}{\sum_j \quad m_j} = \frac{\sqrt{(\sum_j \quad m_j cos\theta_j)^2 + (\sum_j \quad m_j sin\theta_j)^2}}{\sum_j \quad m_j}, \qquad \theta_1 = 0,\ \theta_j = \sum_{k<j} \quad a_k, \qquad (S6)$$

where $a_k$ is the sector angle between junctions $k$ and $k+1$. At mechanical equilibrium, $I = 0$.

We find that across the stationary vertices, the median CV is about 12%, and the median vector imbalance is 0.14, with most vertices between 0.05 and 0.15 (Fig. S2E–F). Both measures are significantly larger at remodeling vertices measured in the same zipping events, as expected for a vertex that is actively driven out of force balance. We don't expect these measures to exactly reach zero at a truly force-balanced vertex, because Myosin II intensity is an indirect proxy for tension. Furthermore, the measured sector angles at a vertex do not exactly sum to $360°$ (mean deviation of order $8°$), and the junctions need not lie in a single plane. Two stationary vertices show larger imbalance (0.22 and 0.31) together with the largest CV. For the majority of stationary vertices, the junctional tensions are close to force balance. The immobility of the stationary vertex is therefore consistent with the force-balance framework and does not require a friction asymmetry.

**Clutch hypothesis –** Force balance constrains the net felt force, $\alpha_2 F_2$, not the coupling $\alpha_2$ on its own. From Eq. (S2), $v_2 \approx 0$ implies only

$$\alpha_2 F_2 \approx 0 \quad \Leftrightarrow \quad (\alpha_2 \approx 0)\ or\ (F_2 \approx 0). \qquad (S7)$$

The Lami analysis shows that $F_2$ is close to zero. A vertex at mechanical equilibrium does not move whatever its coupling, so its immobility cannot discriminate an engaged clutch from a disengaged one. As we show with the simulation and experiments, the endogenous calcium flashes are confined to the receding cell and do not reach the stationary vertex, so that vertex is never driven, and its lack of motion is uninformative about $\alpha_2$.

The remodeling vertex is where the clutch hypothesis is non-vacuous. It carries an active pull ($F_1 > 0$, the Myosin II horseshoe) and it moves ($v_1 > 0$), which by (S2) requires $\alpha_1 > 0$. The molecular data are consistent with an engaged transmission pathway at this vertex: ZO-1, an adaptor linking tight-junction transmembrane proteins to actomyosin, is enriched there (Fig. 3K–L), and the TCJ protein Angulin-1 is comparatively stable there (Fig. 3J), whereas the stationary vertex has lower ZO-1 and a rapidly exchanging Angulin-1 population. The product of Myosin II and ZO-1 intensities at the remodeling vertex, a combined measure of contractility and coupling, shows local increases that precede accelerated elongation (Fig. S2L–M), although its instantaneous level does not predict elongation rate. These observations establish differential composition consistent with differential force transmission; they do not by themselves establish that the stationary vertex would fail to move if force were applied to it. A direct test is to deliver a calcium flash at the stationary vertex.

**Calcium-gated dynamics and the response to an induced flash** – The force-balance relation in Eq. (S2) treats $\alpha_i$ and $\lambda_i$ as fixed. The calcium data show that contractility is pulsatile. Calcium flashes in the extension precede F-actin accumulation by roughly 13 s (Fig. 5E–F), Myosin II is comparatively steady while F-actin pulses (Fig. 4A, F, 5C–D), ZO-1 at the remodeling vertex rises after an induced flash (Fig. 6E–F), and the rate of BCJ elongation increases after both endogenous and induced flashes (Fig. 5H, 6G). In the model, we therefore let calcium drive the F-actin arm of the contractile machinery and let engagement follow contractility.

Denoting $c(t)$ as the calcium signal, we have,

$$\tau_f \dot{f} = -(f - f_0) + \beta\, c(t), \qquad (S8)$$

$$a(t) = m\, f(t), \qquad (S9)$$

$$\alpha(t) = \chi\, s\big(a(t)\big), \qquad s(a) = \frac{a^n}{a^n + a_*^n}, \qquad (S10)$$

$$\lambda_1(t) = \lambda_0 + \kappa\, a(t), \qquad F(t) = 2\lambda_1(t) cos\theta_1 - \lambda, \qquad (S11)$$

$$\gamma_i \; = [\alpha(t)\, F(t)]_{+}. \qquad (S12)$$

Here $f$ is F-actin activity, which relaxes to a baseline $f_0$ with time constant $\tau_f$ and is driven by calcium with gain $\beta$, $m$ is the steady local Myosin II level, $a = mf$ is the contractile activity, and $\chi \in [0,1]$ is the force transmission coefficient that is set by the junctional composition (ZO-1, Angulin-1). The function $s(a)$ is a saturating clutch engagement switch with threshold $a_*$, so that a slack network transmits little force, and $l$ is the length of the bicellular junction.

The bracket in Eq. (S12) denotes the positive part, $[x]_+ = max(x, 0)$, which implies rectification. Equation (S12) therefore gives $\gamma\dot{l} = \alpha F$ when the transmitted force pulls the remodeling vertex forward ($\alpha F > 0$) and $\dot{l} = 0$ when it would pull the vertex back ($\alpha F \le 0$). For the parameters of Table S1 the rectification is never active. The net force at baseline contractility is $F(a_0) = 2(\lambda_0 + \kappa a_0)cos\theta_1 - \lambda = 0.93$, and since $\lambda_1 \ge \lambda_0$ for any $a \ge 0$. Therefore, $F \ge 2\lambda_0 cos\theta_1 - \lambda = 0.03 > 0$ throughout every simulation in Fig. 6, where (S12) reduces to $\gamma\dot{l} = \alpha F$. Averaging (S12) over many flashes recovers Eq. (S2) with $\alpha_1 = \chi\langle s(a)\rangle$, i.e. the force-balanced quasi-static model is the pulse-averaged limit of the dynamic one.

Endogenous flashes are modeled as a train of symmetric pulses, $c(t) = \sum_k A_k \exp[-(t - t_k)^2/(2\sigma_c^2)]$ith random timescales $t_k$. The bicellular junction then elongates continuously, with a rate $\dot{l}$ that rises after each flash and relaxes between flashes. The asymmetry between the two vertices is due to differences in $\chi$ and the spatial confinement of $c(t)$ to the remodeling vertex, not by the instantaneous value of any single signal. This is why velocity does not correlate with the instantaneous product of Myosin II and ZO-1 (Fig. S2L-M), while local increases in that product precede acceleration.

A calcium-uncaging event is modeled as an instantaneous rise at $t = 0$ followed by a two-component decay,

$$c(t) = A\,\Theta(t)\big[w\, e^{-t/\tau_{fast}} + (1 - w)\, e^{-t/\tau_{slow}}\big], \qquad (S13)$$

matching the shape of the measured transient (Fig. 6B). The slow decay time $\tau_{slow}$ is the burst duration, which is variable between experiments (Fig. 6A). Eqs. (S8)–(S12) are integrated from $t = \; -2$ to $+2$ min, and the slope of $l(t)$ is fitted over the 2 min before and after the flash, as for the data. Fig. 6I shows the cascade for a single flash: calcium, then F-actin production with delay $\tau_f$, then engagement $\alpha$ (related to the ZO-1 rise in Fig. 6E–F), then a sustained increase in elongation rate. Fig. 6J shows pre- and post-flash slopes for ten simulated flashes with amplitude and duration drawn from the ranges seen experimentally (Fig. 6G). Fig. 6K shows the difference between post-flash and pre-flash slopes (i.e., difference in elongation rate) as a function of burst duration for three amplitudes. Short or weak bursts that do not carry $a$ across the engagement threshold produce little acceleration. The response saturates for long bursts, consistent with the weak correlation between F-actin increase and burst duration (Fig. S4E).

**Table 1. Parameters of the calcium-gated model used for Fig. 6H-K. The parameter values are illustrative and are not fitted.**

| Symbol | Meaning | Value (Fig. 6) | Basis |
|---|---|---|---|
| $\tau_f$ | F-actin response time | 0.45 min | F-actin peak 0.5–1 min post-flash (Fig. 6D) |
| $f_0$ | baseline F-actin activity | 1 | normalization |
| $\beta$ | calcium-to-F-actin gain | 1.5 | peak F-actin $\approx 1.5 \times$ baseline (Fig. 6D) |
| $m$ | steady Myosin II (remodeling vertex) | 1 | normalization |
| $\chi$ | clutch competence | 1 | remodeling vertex engaged |
| $n$, $a_*$ | engagement switch | 3, 1.8 | baseline creep $\ll$ post-flash rate |
| $\lambda_0$, $\kappa$ | shoulder tension, baseline and gain | 0.50, 0.60 | dimensionless |
| $\lambda$, $cos\theta_1$ | Bicellular junction tension, geometry | 0.72, 0.75 | acute remodeling-vertex angle |
| $\gamma$ | friction coefficient | 1 | velocity scale 0.1–1 $\mu$m/min |
| $A$, $w$ | flash amplitude, fast fraction | 1 (0.5–1.5), 0.5 | Fig. 6B |
| $\tau_{fast}$, $\tau_{slow}$ | flash decay times | 0.12 min, 1.5 min (0.15–4) | Fig. 6B, S4D |

**Video Legends**

**Video S1: The *Xenopus laevis* embryonic epithelium becomes more regularly packed during the blastula-to-gastrula transition, related to Fig. 1A.**

Live imaging of a *Xenopus laevis* embryo expressing membrane probe (BFP-mem, mpl-inferno LUT) starting at late stage 8 (blastula). Time shown in hr:min. Scale bar, 50 μm. Time interval, 5 min; video frame rate, 5 fps.

**Video S2: Tricellular Zipping: a cell-cell junction remodeling process that resolves long, thin cell extensions, related to Fig. 1C.**

Live imaging of a *Xenopus laevis* embryo mosaically expressing two different membrane probes: BFP-mem and mCherry-farnesyl. During tricellular zipping, a long, thin cell extension is zipped up via unidirectional movement of the remodeling tricellular vertex (green arrow), while the opposing vertex remains stationary (gray arrow), leading to the formation of a *de novo* bicellular interface. Time shown in min:sec. Scale bar, 20 μm. Time interval, 78.90 seconds; video frame rate, 5 fps.

**Video S3: Tricellular zipping resolves cellular extensions that arise during cell divisions, related to Fig. 2A.**

Live imaging of a *Xenopus laevis* embryo expressing a membrane probe (BFP-mem) and a F-actin probe (LifeAct-miRFP703). Video illustrates that neighboring cells are pulled inward during cell divisions creating long, thin extensions (white box). These extensions are resolved by tricellular zipping, marked by the unidirectional movement of the remodeling vertex (green arrow). Time shown in hr:min. Scale bar, 50 μm. Time interval, 5 min (video pauses briefly to highlight the long cell extension); video frame rate, 5 fps.

**Video S4: Tricellular tight junction protein Angulin-1 accumulates at the remodeling vertex and FRAP of Angulin-1 at the remodeling and stationary vertices, related to Fig. 3H-J.**

The first video shows Angulin-1-3xGFP (Sepia LUT) accumulates at the remodeling vertex (yellow arrow) during tricellular zipping. Time shown in min:sec. Time interval, 31.67 seconds; video frame rate, 5 fps. The second video shows Angulin-1-3xGFP (FIRE LUT) at remodeling and stationary vertices before, during, and after photobleaching. Bleaching is indicated by white circles. Images are shown using the FIRE LUT, adjusted in the same way for both videos. Time shown in seconds. Scale bars, 10 μm; Time interval, 0.88 seconds; video frame rate, 20 fps.

**Video S5: Tight junction protein ZO-1 dynamically accumulates at the remodeling vertex during tricellular zipping, related to Fig. 3K.**

Live imaging of a *Xenopus laevis* embryo expressing ZO-1 (BFP-ZO-1, inverted gray LUT). Video illustrates that the tight junction protein ZO-1 accumulates at the remodeling vertex (green arrow) during tricellular zipping and is highly dynamic at the zipping front (magenta arrow). Time shown in min:sec. Scale bar, 10 µm. Time interval, 27.23 seconds; video frame rate, 5 fps.

**Video S6: Myosin II dynamics during tricellular zipping, related to Fig. 4A.**

Live super-resolution imaging of a *Xenopus laevis* embryo expressing a probe for Myosin II (SF9-mCherry, FIRE LUT). Video illustrates that Myosin II accumulates more at the remodeling vertex (green arrow) than at the stationary vertex (gray arrow) at the beginning of tricellular zipping. As zipping progresses, a horseshoe-like accumulation of Myosin II is observed at the zipping front and moves together with the remodeling vertex. The horseshoe-like accumulation dissipates as the remodeling vertex stops moving. Time shown in min:sec. Scale bar, 10 µm. Time interval, 27.36 seconds (early-stage video) and 29.36 (late-stage video); video frame rate, 5 fps.

**Video S7: Pulsatile F-actin dynamics within the receding cell extension during tricellular zipping, related to Fig. 4F.**

Live super-resolution imaging of a *Xenopus laevis* embryo mosaically expressing two different F-actin probes: LifeAct-GFP and LifeAct-RFP. Video illustrates pulsatile F-actin dynamics within the long cell extension during tricellular zipping. Yellow arrow highlights repeated increase in F-actin signal at the zipping front in the receding cell. Time shown in min:sec. Scale bar, 10 µm. Time interval, 165.12 seconds; video frame rate, 2 fps.

**Video S8: Localized calcium flashes within the long cell extension during tricellular zipping, related to Fig. 5A.**

Live imaging of a *Xenopus laevis* embryo expressing a probe for F-actin (LifeAct-miRFP703) and a cytoplasmic calcium indicator (GCaMP6m). Video illustrates F-actin and calcium dynamics within the long cell extension during tricellular zipping. Yellow arrow highlights repeated flashes of calcium associated with zipping. Time shown in min:sec. Scale bar, 20 µm. Time interval, 22.23 seconds; video frame rate, 5 fps.

**Video S9: Induced calcium bursts promote rapid increase in speed of tricellular zipping, related to Fig. 6A.**

Live imaging of a *Xenopus laevis* embryo expressing a cytoplasmic calcium indicator (GCaMP6m; FIRE LUT), ZO-1 (mRFP-ZO-1; inverted gray LUT), and a probe for F-actin (LifeAct-miRFP703; Green Blue Fire LUT) pre- and post- calcium uncaging. Embryos were incubated in DMNP-EDTA (caged calcium), -AM ester and imaged in the dark, followed by a 405-laser stimulation of the zipping junctions, causing a burst of calcium and increased rate of tricellular zipping. Time shown in min:sec. Scale bar, 10 μm. Time interval, 20.17 seconds; video frame rate, 7 fps.

**Video S10: Stochastic single-cell calcium flashes during ventral midline invagination and germband extension in *Drosophila* embryo, related to Fig. S5A.**

Live imaging of a *Drosophila melanogaster* embryo expressing a membrane-bound probe for cytosolic calcium (myr-GCaMP8s). Video illustrates that calcium transiently increases in individual cells. Time shown in min:sec. Scale bar, 15 μm. Time interval, 7.588 seconds; video frame rate, 7.6 fps.

**Video S11: Ionomycin injection at early stage 5 prevented apical constriction of the mesoderm, injection at stage 6 resulted in failed internalization of the mesoderm in *Drosophila* embryo, related to Fig. 7C.**

Live imaging of a *Drosophila melanogaster* embryo expressing membrane-bound calcium probe GCaMP8s (myr-GCaMP8s) after 1 mM Ionomycin injection at Stage 5 and Stage 6. Time shown in min: sec. Scale bar, 10 μm. Time interval, 39.11 seconds; video frame rate, 6 fps.
